# TopBP1 biomolecular condensates: a new therapeutic target in advanced-stage colorectal cancer

**DOI:** 10.1101/2024.09.10.612204

**Authors:** Laura Morano, Nadia Vezzio-Vié, Adam Aissanou, Tom Egger, Antoine Aze, Solène Fiachetti, Benoît Bordignon, Cédric Hassen-Khodja, Hervé Seitz, Louis-Antoine Milazzo, Véronique Garambois, Laurent Chaloin, Nathalie Bonnefoy, Céline Gongora, Angelos Constantinou, Jihane Basbous

**Affiliations:** Institut de Génétique Humaine, Univ Montpellier, CNRS, Montpellier, France; IRCM, Univ Montpellier, ICM, INSERM, Montpellier, France; Institut de Recherche en Infectiologie de Montpellier (IRIM), Univ Montpellier, CNRS, Montpellier, France; Montpellier Ressources Imagerie, BioCampus, University of Montpellier, CNRS, INSERM, Montpellier, France

**Author notes:** Deceased (Angelos Constantinou).

## Abstract

In cancer cells, ATR signaling is crucial to tolerate the intrinsically high damage levels that normally block replication fork progression. Assembly of TopBP1, a multifunctional scaffolding protein, into condensates is required to amplify ATR kinase activity to the levels needed to coordinate the DNA damage response and manage DNA replication stress. Many ATR inhibitors are tested for cancer treatment in clinical trials, but their overall effectiveness is often compromised by the emergence of resistance and toxicities. In this proof-of-concept study, we propose to disrupt the ATR pathway by targeting TopBP1 condensation. First, we screened a molecule-based library using a previously developed optogenetic approach and identified several TopBP1 condensation inhibitors. Amongst them, AZD2858 disrupted TopBP1 assembly induced by the clinically relevant topoisomerase I inhibitor SN-38, thereby inhibiting the ATR/Chk1 signaling pathway. We found that AZD2858 exerted its effects by disrupting TopBP1 self-interaction and binding to ATR in mammalian cells, and by increasing its chromatin recruitment n cell-free *Xenopus laevis* egg extracts. Moreover, AZD2858 prevented S-phase checkpoint induction by SN-38, leading to increased DNA damage and apoptosis in a colorectal cancer cell line. Lastly, AZD2858 showed synergistic effect in combination with the FOLFIRI chemotherapy regimen in a spheroid model of colorectal cancer.

**Graphical abstract:** 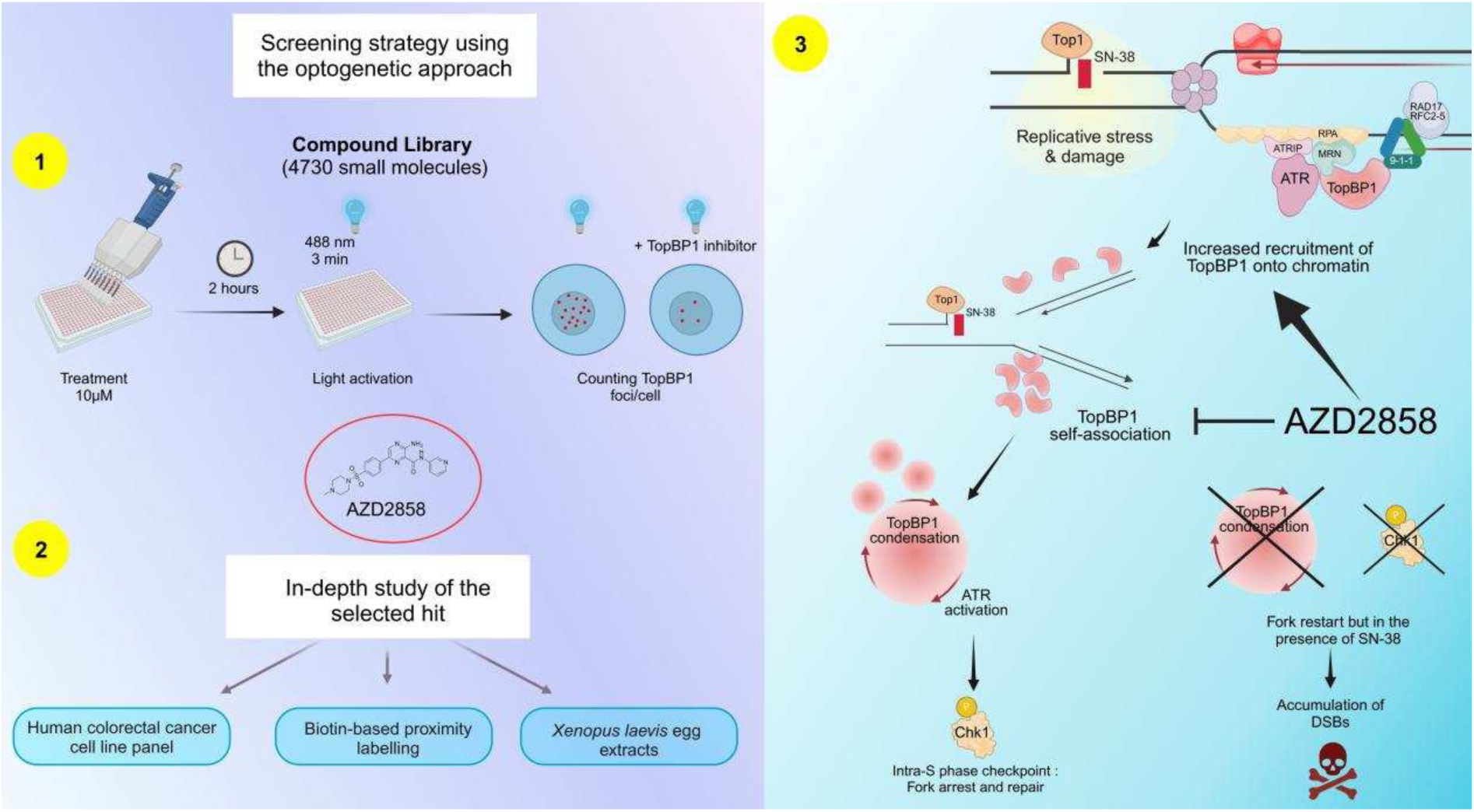

## Introduction

In every cell, thousands of DNA lesions are induced by endogenous and environmental agents (1). To overcome their deleterious effects, cells have evolved an intricate network of DNA damage response pathways that detect, signal and repair DNA lesions. These pathways work in coordination with key physiological processes, such as cell cycle progression, chromatin remodeling, and transcription (2,3). Cancer cells typically exhibit higher levels of genotoxic stress than non-transformed cells due to their altered metabolism and high proliferation rate (4–6). As a complement to radiotherapy and chemotherapy that overload cancer cells with DNA lesions, the aim of novel anti-cancer strategies is to increase the sensitivity of cancer cells to genotoxic stress by targeting DNA damage signaling and DNA repair mechanisms (4,7).

For example, ATR signaling activates cell cycle checkpoints (8–10), promotes DNA repair (11), regulates the firing of replication origins (12), and the cellular pool of deoxyribonucleotides (13–16). The ATR signaling pathway is also essential for the proliferation of cancer cells that display intrinsically higher levels of genotoxic stress. Therefore, several clinical trials are evaluating the efficacy of ATR inhibitors in patients with cancer (17), particularly cancers in which the oncogene CDC25A is overexpressed (18). However, as kinase inhibitors often exert a strong selective pressure for the acquisition of drug resistance through kinase mutations, alternative therapeutic options must be identified (19).

Unlike classical approaches that target the activity of a specific protein with small molecule inhibitors, recent cell biology advances suggest that targeting biomolecular condensates may provide conceptually novel approaches to drug discovery (20). Biomolecular condensates are subcellular compartments that selectively concentrate hundreds of proteins and nucleic acids, without a surrounding membrane (21,22). These structures underlie the spatiotemporal organization of multiple biological processes. Moreover, the aberrant properties of condensates have been implicated in many diseases, including neurodegeneration, cardiomyopathy, cancer and viral infections (20). Condensate formation is typically driven by multivalent scaffold proteins that function as central nodes in molecular networks (21,22). Therefore, one potential approach to disturb the condensate properties and functions is to interfere with their essential protein-protein or protein-nucleic acid interactions.

In humans, the main ATR activator in S phase is Topoisomerase IIβ binding protein 1 (TopBP1), a scaffold protein that includes nine BRCA1 carboxyl terminal (BRCT) protein-protein interaction motifs (23,24) and an intrinsically disordered ATR-activation domain located between BRCT6 and BRCT7-8 (23,25). Recent studies indicate that TopBP1 activates the ATR signaling pathway via the formation of biomolecular condensates, which appear as nuclear foci by immunofluorescence microscopy (26). The available evidence suggests that ATR activation is a multistep process (27,28). TopBP1 is recruited to ATR-activating DNA structures by MRE11 (29) and binds to RPA (30), ATRIP (31) and the 9-1-1 complex (32,33) to form a stable ATR-activating complex. Then, TopBP1 is phosphorylated by the basal kinase activity of ATR, triggering the assembly of TopBP1 condensates (26). We found that TopBP1 condensation functions as a molecular switch that amplifies ATR activity up to the level required for signal transduction by the checkpoint effector kinase Chk1 (26).

Here, we explored the feasibility of targeting TopBP1 condensates as a novel strategy to interfere with ATR signaling and sensitize colorectal cancer (CRC) cells to chemotherapeutic drugs. We present a high throughput optogenetic screening approach to identify small-molecule modulators of TopBP1 condensation. Using this approach, we identified AZD2858, a known glycogen synthase 3β (GSK-3β) inhibitor, as a potential modulator of TopBP1 condensation. We demonstrated in a CRC cell line that AZD2858 (at very low doses) inhibits the assembly of endogenous TopBP1 and suppresses the activation of the ATR/Chk1 pathway induced by the topoisomerase I inhibitor SN-38, the active metabolite of irinotecan. Furthermore, the combination of AZD2858 with FOLFIRI (5-fluorouracil + SN-38) displayed a synergistic effect in CRC cell spheroid cultures, including in spheroids from CRC cells resistant to SN-38. Incubation with AZD2858 for 6-12 hours improved the induction of apoptosis by SN-38 and FOLFIRI by dampening the S checkpoint. Our observations suggest that the cytotoxic effects observed with the AZD2858 and SN-38 combination are mainly explained by disruption of TopBP1 assembly, rather than by the direct involvement of the GSK-3β pathway. These data support the feasibility of targeting condensates formed in response to DNA damage to improve chemotherapy-based cancer treatments.

## Materials and methods

### Cell culture

Flp-In# 293 T-REx cells were obtained from Thermo Fisher and cultured in Dulbecco’s Modified Eagle Medium (DMEM)-GlutaMAX medium (Merck-Sigma-Aldrich, #D5796) supplemented with 10% heat-inactivated fetal bovine serum (FBS). For the generation of stable cell lines that express TopBP1 fused to the photoreceptor cryptochrome 2 (Cry2) and mCherry (optoTopBP1), please refer to Frattini et al., 2024 (26). The HCT116 human CRC cell line was obtained from Horizon (#HD-PAR-082). The human SW620, SW480, HT29 and murine CT26 CRC cell lines from ATCC. (CT26.WT -CRL-2638, HCT116-CCL247,

HT29-HTB-38, SW620-CCL-227, SW480-CCL228) The control human colon cell line CCD 841 CoN was also obtained from ATCC (CCD 841 CoN-CRL-1790). The SN-38-resistant clones HCT116-SN6 (low-level resistance) and HCT116-SN50 (high-level resistance) were generated in the laboratory as previously described (34). Briefly, parental HCT116 cells were grown in the presence of 10 nmol/L or 15 nmol/L SN-38, respectively. SW620 shLuc and SW620 shGSK3-β cells were kindly provided by Maguy Del Rio. The description of their generation can be found in Cherradi et al. 2023 (35). All CRC cell lines were grown in and cultured in RPMI medium (Merck-Sigma-Aldrich, #R8758) supplemented with 10% heat-inactivated FBS. All cell lines were maintained at 37°C in saturated humidity with 5% CO_2_. All cell lines were tested and authenticated by short-tandem repeat profiling (Eurofins Genomics). Cells were routinely tested for mycoplasma contamination.

### Drugs and antibodies

A personalized TargetMol library containing 4730 molecules was used for the high throughput screening (see Screening section). SN-38 (active metabolite of irinotecan, Tocris #2684) was diluted to 10 mM in DMSO and stored at -20°C. 5-fluorouracil (5-FU; Sigma, #F6627) was diluted to 10 mM in water and stored at room temperature (RT). AZD2858 (Euromedex, #AB-M2178) was diluted to 10 mM in DMSO and stored at -80°C. The drugs were diluted in the adapted medium and tested at the concentrations indicated in the figures and legends.

The following primary and secondary antibodies were used are indicated in Table S6

### Screening

OptoTopBP1-expressing Flp-In 293 T-Rex cells were seeded in a 384-well plate at a density of 10,000 cells/well in DMEM supplemented with 10% FBS and 2 µg/ml doxycycline and allowed to attittiach overnight. Then, cells were incubated with the TargetMol library compounds at the concentration of 10 µM for 2 h. From the 10 mM stock solutions in DMSO, each compound was diluted to 2 mM (intermediated dilution) before addition to the wells by automated pipetting at a final concentration of 10µM. After incubation, cells were exposed to cycling pulses (4 s 8ON9, 10 s 8OFF9) of 488 nm blue light for 5 min. Cells were fixed immediately in 4% paraformaldehyde (PFA) (Euromedex, #15710-S) diluted in phosphate-buffered saline (PBS) for 15 min. Cells were rinsed with PBS twice and permeabilized in PBS/0.2% Triton X-100 for 10 min. Cells were rinsed in PBS twice and nuclei were counterstained with 1 µg/mL Hoechst 33342 (Thermo Fisher Scientific, #62249). Imaging was performed with the Opera PHENIX system at the Montpellier Ressources Imagerie facility.

### Western blotting

Whole cell extracts were obtained by lysing cells in RIPA buffer (50 mM Tris, 150 mM NaCl, 1% NP-40, 1% deoxycholate, 0.1% SDS, pH 8) on ice for 30 min. After sonication (40% amplitude, 3 cycles of 3 s sonication and 3 s resting), the protein amount was quantified using the Pierce# BCA Protein Assay Kit (Thermo Fisher #23225). Laemmli buffer was added, and protein extracts were boiled at 95°C for 5 min. 40 μg of protein samples were resolved on precast SDS-PAGE gels (4-15%, 7.5% and 10%, BioRad) and transferred to nitrocellulose membranes using the Bio-Rad Trans-Blot Turbo transfer device. Membranes were saturated in 5% non-fat milk diluted in TBS (20 mM Tris, 150 mM NaCl, pH 7.4)-0.1 % Tween 20, and incubated with primary antibodies at 4°C overnight. The used antibodies were against Chk1 phosphorylated at S345 (Cell Signaling Technology, #2348), Chk1 (Santa-Cruz Biotechnology, #sc8408), Chk2 phosphorylated at T68 (Cell Signaling Technology, #2661S), Chk2 (Milipore, #05-649), TopBP1 (Euromedex, #A300-111A or Santa Cruz Biotechnology, #sc-271043), ATM phosphorylated at S1981 (Rockland, #200-301-400), α-tubulin (Sigma, #T5168), RPA phosphorylated at S33 (Abcam, #ab2118877), RPA (Abcam, #AB2175), GSK-3β phosphorylated at S9 (Cell Signaling Technology, #9336), GSK-3β (Cell Signaling Technology, #9832), PARP1 (Santa Cruz Biotechnology, #sc-8007), cleaved caspase-3 (Cell Signaling Technology, #9661). For more information, please refer to **Table S6**. Membranes were then incubated with anti-mouse (Cell Signaling Technology, #7076S) or anti-rabbit HRP (Cell Signaling Technology, #7074S) secondary antibodies for 1 h. Revelation was carried out with ECL Clarity (BIO-RAD, #170-5061) or ECL select (BIO-RAD, #1705062) according to the manufacturer9s instructions.

### Immunofluorescence analysis

Cells were grown on coverslips and after incubation with the corresponding drugs (see above) for 2 h, the soluble fraction of cells was removed during the pre-extraction step in Cytoskeleton buffer <CSK= (PBS containing 0.2% Triton X-100) at 4°C for 60 s. Cells were then fixed with 4% PFA/PBS at RT for 20 min. Non-specific epitopes were saturated with a blocking solution (5% BSA/PBS) at RT for 30 min. For immunostaining, anti-TopBP1 (Santa Cruz Biotechnology, #sc-271043, diluted at 1/100), anti-53BP1 (Cell Signaling Technology, #4937S) or anti-PML (Santa Cruz Biotechnology, #sc-966, diluted at 1/300) primary antibodies and the anti-mouse coupled to fluorochrome (Invitrogen, #A-11011, diluted at 1/500) secondary antibody were diluted in blocking solution and incubated for 1 h or 45 min, respectively. Three 5-min washes between incubation steps were performed in PBS/0.1% Tween 20 at RT. Hoechst was used at 1 µg/mL for DNA staining and coverslips were mounted on slides with the Prolong Gold antifade reagent (Invitrogen, #P36930). Images were captured using a 63 X objective and a Zeiss AxioImager with ApoTome at the Montpellier Ressources Imagerie facility.

### TurboID assay: pulldown of biotinylated proteins

Flp-In 293 T-Rex cells that express optoTopBP1 and grown to 75% of confluence were incubated with 2 µg/ml of doxycycline for 16 h. The next day, during the last 15 min of incubation with the indicated drugs, 500 mM of biotin was added. Cells were then washed with PBS and lysed in lysis buffer (50 mM Tris-HCl pH 7.5, 150 mM NaCl, 1 mM EDTA, 1 mM EGTA, 1% NP-40, 0.2% SDS, 0.5% sodium deoxycholate) supplemented with 1X complete protease inhibitor, 1X phosphatase inhibitor and 250 U benzonase. Lysed cells were incubated on a rotating wheel at 4°C for 1 h before sonication (40% amplitude, 3 cycles of 1 s sonication – 2 s resting) on ice. After centrifugation at 7750 rcf at 4°C for 30 min, cleared supernatants were transferred to new tubes and the total protein concentration was determined using the Bradford protein assay (BioRad). For each condition, 1 mg of proteins was incubated with 100 µl of streptavidin-agarose beads on a rotating wheel at 4°C for 3 h. After centrifugation at 400 rcf for 1 min, beads were washed sequentially with 1 ml lysis buffer, 1ml wash buffer 1 (2% SDS in H_2_O), 1 ml wash buffer 2 (0.2% sodium deoxycholate, 1% Triton X-100, 500 mM NaCl, 1 mM EDTA, and 50 mM HEPES, pH 7.5), 1 ml wash buffer 3 (250 mM LiCl, 0.5% NP-40, 0.5% sodium deoxycholate, 1 mM EDTA, 500 mM NaCl and 10 mM Tris pH 8) and 1 ml wash buffer 4 (50 mM Tris pH 7.5 and 50 mM NaCl). Bound proteins were eluted from the agarose beads with 80 µl of 2X Laemmli sample buffer and incubated at 95°C for 10 min. Western blot analysis was performed as previously described to analyze the self-proximity of optoTopBP1 molecules.

### *Xenopus laevis* egg extract assay

Interphasic Low Speed Egg extracts (LSE) and demembranated sperm nuclei were prepared as previously described (Menut S et al 1999). LSE were then clarified by centrifugation at 20,000 rpm in an SW55Ti rotor for 20 min. Chromatin preparation was performed as described before (Aze et al; 2016). Briefly, demembranated sperm nuclei were added at a final concentration of 4000 nuclei per microliter to LSE supplemented with energy regeneration mix (200 µg/ml creatine phosphokinase, 200 mM creatin phosphate, 20 mM ATP, 20 mM MgCl_2_, 2 mM EGTA) and cycloheximide (250 µg/ml). When indicated, LSE were pre-incubated with camptothecin (CPT), AZD2858, or DMSO (as control) for 5 min. The mixtures were incubated at 23°C for the indicated times, then samples were diluted in EB buffer (100 mM KCl, 50 mM Hepes-KOH, 2.5 mM MgCl_2_, supplemented with 0.25% NP40) and centrifuged over a 30% sucrose cushion. Purified chromatin pellets were washed once with EB buffer and recovered in Laemmli buffer for western blot analysis. Antibody references are provided in **Table S6**.

### Cell cycle assay

Cell cycle profiling using 5-bromo-2′-deoxyuridine (BrdU)/propidium iodide (PI) was performed as previously described (Egger et al., 2022). Briefly, cells were pulsed with 10 μM BrdU for 15 min and fixed in ice-cold 70% ethanol. Cells were digested in 30 mM HCl, 0.5 mg/mL pepsin (37°C, 20 min). DNA was denatured in 2 N HCl (RT, 20 min) and BrdU was immunodetected using the anti-BrdU clone BU1/75 in PBS/2% goat serum/0.5% HEPES/0.5% Tween 20 and revealed with a goat anti-rat Alexa Fluor 488 secondary antibody. DNA was stained with 25 μg/mL PI in PBS and cells were analyzed on a Gallios flow cytometer (Beckman Coulter) at the Montpellier Ressources Imagerie facility. 20,000 cells/sample were analyzed using the Kaluza software.

### Detection of apoptotic cells with the sub-G1 assay

For visualization of sub-G1 apoptotic cells by flow cytometry, the <SubG1 Analysis Using Propidium Iodide= protocol developed by the UCL - London’s Global University was used. Briefly, cells were collected and fixed in ice-cold 70% ethanol, followed by two washes in phosphate-citrate buffer (192 mM Na_2_HPO_4_, 4 mM citric acid, pH 7.8). Ribonuclease A (100 μg/ml in PBS) digestion was performed at 37°C for 30 min. DNA was stained with PI (50 µg/ml in PBS) for 15 min. Cells were analyzed on a Gallios flow cytometer (Beckman Coulter) at the Montpellier Ressources Imagerie facility. 20,000 cells/sample were analyzed with the Kaluza software. Debris (low FSC/SSC) was kept and the sub-G1 fractions were calculated as the percentage of sub-G1 (<2N PI content) cells relative to the total number of events.

### Pulse Field Gel Electrophoresis (PFGE)

Cells were collected and embedded in 0.5% agarose plugs (in PBS), at the exact concentration of 0.7×10^6^ cells per plug. Plugs were incubated in lysis buffer (100 mM EDTA pH8, 0.2% sodium deoxycholate, 1% sodium lauryl sarcosine, 1 mg/ml proteinase K) at 37°C for 48h. Plugs were washed 3 times in wash buffer (20 mM Tris pH 8, 50 mM EDTA pH 8) and inserted in 0.9% agarose gel prepared in 0.5X TBE (from a 5X stock PanReac-AppliChem #A4228). Chromosomes were separated by pulsed-field gel electrophoresis for 23 h (Biometra Rotaphor 8 System, 23 h; interval: 30-5 s log; angle: 120-110 linear; voltage: 180-120 V log, 13°C). Then, gels were stained with ethidium bromide (0.5 µg/mL) for analysis.

### 2D cell growth inhibition assay

Cell growth was evaluated using the sulforhodamine B (SRB) assay, as described by Skehan et al., 1990 (36). Briefly, 500 cells/well were seeded in 96-well plates. After 24 h, cells were incubated with the indicated drug concentrations for 96 h. Cells were fixed in 10% trichloroacetic acid solution (Sigma, #T9159), rinsed in water and stained with 0.4% SRB (Sigma, #S9012) in 1% acetic acid (Fluka, #33209). Plates were washed three times with 1% acetic acid and fixed SRB was dissolved in 10 mmol/L Tris-Base solution (Trizma®base SIGMA, #T1503). The absorbance was read at 560 nm using a PHERAstar FS plate reader (BMG Labtech). Non-treated controls were used to normalize the cell growth inhibition values. The IC_50_ of drugs were determined graphically using the cell growth inhibition curves.

### 3D spheroid assay

For spheroid generation, 100 µL/well of cell suspensions at optimized densities (50 cells/well) were dispensed in ultralow attittiachment 96-well round-bottittiomed plates (Corning B.V. Life Sciences, #7007) and cultured at 37°C, 5% CO_2_, 95% humidity. Cells were first incubated with the indicated drugs at day 1 and then at day 4. At day 7, images were captured with the Celigo Imaging Cytometer (Nexcelom Bioscience) using the “Tumorosphere” application. Cell viability was measured using a CellTiter-Glo Luminescent Cell Viability Assay (Promega, #G9683), according to the manufacturer’s instructions. Luminescence was measured in the white 96-well plates using a PHERAstar FS plate reader (BMG LABTECH). The IC_50_ was determined graphically from the cytotoxicity curves.

### Synergy matrix

The percentage of living cells after incubation with each drug alone or in combination was calculated and normalized to that of untreated cells. Then, using a script developed by Tosi et al., 2018 (37) in the <R= software based on the effect of each molecule alone (Bliss and Lehàr equation), a synergy matrix was generated. This matrix associates a number to each drug combination: if positive (red), it indicates synergistic effects; if negative (green), it indicates antagonistic effects. Additives effects (∼ 0) are displayed in black.

### In vitro cytotoxicity assays

Cells were seeded at 500 cells/well in black flat-bottittiom 96-well plates (Sarstedt, #83.3924), cultured for 24 h, then incubated with AZD2858 or/and FOLFIRI for 96 h. Then, PI (Sigma, #P4864) and Hoechst 33342 (Thermo Fisher Scientific, #62249) were added to the plates at the final concentrations of 1 and 5 µg/mL, respectively. Cells were incubated at 37°C for 30 min before counting the positive cells for each fluorescence signal using a Celigo Imaging Cytometer (Nexcelom Bioscience) and the “Expression analysis – Cell Viability - Dead + Total” application. The number of living cells was then calculated by subtracting the number of dead, PI-positive cells, from the number of total cells given by the Hoechst staining. The percentages of live and dead cells were plottittied and cytotoxic and cytostatic profiles were determined, as previously described by Vezzio-Vie et al., 2022 (38).

### Celigo imaging cytometer-based immunofluorescence analysis

HCT116 cells were seeded at 1,200 cells per well for 48 h or 24 h in black 384-well plates (Greiner, #781091) and incubated with increasing concentrations of AZD2858 and/or FOLFIRI (diluted to 1/2, corresponding to 50 nM of SN-38 and 6 µM of 5-FU) for 2 and 20 h, respectively. Then, cells were fixed in 4% PFA, permeabilized in PBS/0.1% Triton X100 (Sigma), and saturated in 3% BSA/PBS. The primary antibodies anti-Chk1 phosphorylated at S345 (Cell Signaling Technology, #2348L), anti-Chk2 phosphorylated at T68 (Cell Signaling Technology, #2661S), and anti-ATM phosphorylated at S1981 Santa Cruz Biotechnology, #sc-47739) were diluted in 1% BSA/PBS and cells were incubated at 4°C under stirring overnight. Cells were washed three times with 0.05% Tween 20/PBS (Tween®20 Polysrobate Technical VWR CHEMICALS). The Alexa Fluor 568 goat anti-rabbit or anti-mouse IgG (H + L) (Thermo Fisher Scientific, #A11011 or #A11004 respectively) secondary antibody was added at RT for 45 min. The anti-γH2AX (phosphorylated at S139) (Abcam, #195189) antibody was diluted (1/1000) in 1% BSA/PBS and added to the cells at RT for 45 min. Cells were washed three times and 1 µg/mL of Hoechst 33342 in PBS was used to counterstain nuclei (37°C, 30 min). Fluorescence signals were acquired using the <Expression analysis – Target 1 + Mask= application on the Celigo Imaging Cytometer (Nexcelom). The percentage of positive cells, the mean signal intensity per well, and the total cell count were calculated using the Nexcelom Bioscience Celigo Satellite software.

### DNA fiber assay

1×10^6^ HCT116 cells were seeded in 6-well plates. Cells were allowed to attittiach overnight and then sequentially pulsed with 20 μM 5-iodo-2′-deoxyuridine (IdU) (20 min) and 200 μM 5-chloro-2′-deoxyuridine (CldU) (20 min). Cells were harvested, washed and diluted in ice-cold PBS to 0.5– 1 × 10^6^ cells/mL. 2 μL of cell suspension was pipettittied on the edges of SuperFrost microscopy slides. After drying at RT for 3–5 min, 7 μL of DNA Spreading Buffer (200 mM Tris–HCl pH 7.5, 50 mM EDTA, 0.5% SDS) was mixed with the cell-containing drops. After drying at RT for 2–5 min, slides were manually tilted (15 ° – 30 °) to spread the DNA fibers that were then fixed in a 3:1 methanol/acetic acid solution at RT for 10 min. Slides were washed in H_2_O and DNA was denatured in 2.5 M HCl for 1 h. Slides were washed and blocked in PBS/1% BSA/0.1% Tween 20 at RT for 1 h. Then, slides were incubated with a mix of anti-IdU (BD Biosciences, Cat# 347580; RRID:AB_10015219) and anti-CldU (Bio-Rad, Cat# OBT-0030, RRID:AB_2314029) antibodies, both diluted (1/25) in PBS/0.1% Tween 20, at 37 °C in saturated humidity for 45 min. Slides were washed in PBS/0.1% Tween 20 five times (2 min/each) and incubated with a mix of secondary antibodies, diluted 1/50 in PBS/0.1% Tween 20, at 37°C for 30 min. Slides were rinsed in PBS/0.1% Tween 20 five times (2 min/each), dried and mounted with 24 × 50 mm coverslips in 30 μL of ProLong Gold Antifade. Twenty representative pictures for each condition were saved on a Zeiss Axio Imager (40 X objective). The CldU tract lengths of individual fibers consisting of a sequence of IdU and CldU tracts were measured using the open source FiberQ software (39). Experiments were repeated three times and the pooled data from three replicates were plottittied using the online tool SuperPlotsOfData (40). The non-parametric unpaired Mann–Whitney test was used to determine the significance of the results.

### Softtiware tools

Immunofluorescence data were acquired with a Zeiss AXIOIMAGER microscope (Zeiss) and the Zen toolkit connect software. For foci quantification, the pipelines generated on CellProfiler 4.2.1 and described in Egger *et. al*, 2024 were used. For western blot images acquisition, ChemiDoc Imaging System (Biorad) and ImageLab 6.1 were used. The R studio software was used to generate the synergy and viability matrixes with a predefined script. GraphPad version 8 was used for graphical representation and statistical analyses. R was used for statistical analyses. The DNA fibers were quantified using open source FiberQ software. For analysis of TopBP1 foci in parental and SN-38-resistant HCT116 cells, and for DNA fiber assay, biological replicates were pooled and plottittied using SuperPlotsOfData (https://huygens.science.uva.nl/SuperPlotsOfData/) (41). Data distribution was displayed as half-violin plots on the right-hand side of the dot plots to help data visualization.

### Statistical analyses

For the immunofluorescence quantification of foci (TopBP1 **Figure 2A** and 53BP1 **Figure S2A**), all conditions were first compared using one-way ANOVA and confirmed using non-parametric unpaired Mann-Whitney tests on GraphPad Prism version 10.2.1 (the non-normal distributions of foci limited the precision of the ANOVA analyses). SN-38 vs SN-38+AZD2858 (Mann-Whitney); TopBP1 : p-value<0.0001 ; 53BP1 : p-value<0.0001. In addition, we conducted in-depth statistical analyses on R to complete the results obtained with GraphPad. Specifically, for TopBP1 foci, an ANOVA analysis of a generalized linear model (modeling TopBP1 foci number with a Poisson distribution) was performed to confirm this. The ANOVA test shows that SN38 and AZD2858 treatments exhibit significantly negative interaction on TopBP1 foci number (SN38 treatment p-value<2.2e-16; AZD2858 treatment p-value<2.2e-16; SN38:AZD2858 interaction p-value<2.2e-16). For PML foci (**Figure S2B**), all conditions were first compared using Kruskal-Wallis on GraphPad Prism and p-value<0.0001 was obtained. Then, Dunn9s multiple comparison test was conducted to specifically compare the Arsenic condition with the following treatments: untreated cells, AZD2858, SN-38, and SN-38+AZD2858. In each comparison, the p-value was <0.0001.

For DNA fiber analyses, groups were first compared using one-way ANOVA and confirmed using unpaired t-tests on GraphPad Prism version 10.2.1 (the non-normal distributions of CldU tracts length limited the precision of the ANOVA analyses, **Figure 3B**). In addition, we conducted in-depth statistical analyses on R to complete the results obtained with GraphPad : ANOVA analysis of a generalized linear model (modeling replication tract length with a gamma distribution) shows that both drugs, as well as their interaction, have a significant effect on tract length (p-value for SN38 treatment: < 2.2e-16; for AZD2858 treatment: 0.01129; for their interaction: 2.651e-06). We did not detect a significant effect of biological replicate identity (p-value=0.26274).

For cytometry analyses, linear regression analysis was performed on R (**Figure 4B**). For **Figure 3C** and **Figure S5**, we performed an ANOVA analysis to evaluate the effect of chase duration (6 h vs. 12 h) and treatment (untreated vs. AZD2858 vs. SN38 vs. AZD2858+SN38) on the proportion of cells in S phase (Endpoint, BrdU positive cells), S-phase (Pulse-chase, BrdU negative cells, between 2N and 4N, green squares) and G2 phase (Pulse-chase, BrdU positive cells, 4N, red squares). For S-phase analysis, both treatment duration (6h vs. 12h) and nature of treatment (untreated, AZD2858-treated, SN38-treated or (AZD2858+SN38)-treated) have a significant effect on the proportion of cells in S phase in the endpoint experiment (p=0.00308 for treatment duration, p=2.14e-06 for nature of treatment). Interaction between treatment nature and duration did not appear significant (p=0.05942): AZD2858-treated and untreated cells tend to have higher S phase cell counts than SN38-treated and (AZD2858+SN38)-treated cells, regardless of treatment duration; and increasing treatment duration (from 6 to 12h) decreases S phase cell count, regardless of treatment nature. For S-entry analysis, both treatment duration (6h vs. 12h) and nature of treatment (untreated, AZD2858-treated, SN38-treated or (AZD2858+SN38)-treated) have a significant effect on the proportion of cells in SE phase in the pulse-chase experiment (p=0.0173 for treatment duration, p=4.03e-06 for nature of treatment). Interaction between treatment nature and duration also has a significant effect (p=0.00865): AZD2858-treated and untreated cells tend to have higher SE cell counts than SN38-treated and (AZD2858+SN38)-treated cells, but that difference is larger after a 6h treatment than a 12h treatment. For G2 phase analysis, treatment duration (6h vs. 12h) has a significant effect on the proportion of cells in G2 phase in the pulse-chase experiment (p=0.000363). The nature of the administered treatment significantly alters the effect of duration (interaction between treatment nature and treatment duration: p=0.000340), with SN38-treated and (AZD2858+SN38)-treated cells exhibiting similar proportion of G2 cells after 6 and 12h of treatment, while untreated and AZD2858-treated cells undergo a drop in G2 cell count between 6 and 12h of treatment. We confirmed these analysis with conventional t-test on Graphpad Prism : S-phase (6h, 12h) p-value< 0.01; S-entry (12h) p-value< 0.01; G2-phase (12h) p=0.0654 (>0.02, ns: non-significant).

In all cases, significance thresholds were as follows: ns: non-significant; ∗: p < 0.05; ∗∗: p < 0.01; ∗∗∗: p < 0.001; ∗∗∗∗: p < 0.0001. Detailed R scripts and data files for the analysis of data shown in **Figures 2A**, **2C**, **3B**, **4B, S1, S3 and S5** are available at https://github.com/HKeyHKey/Morano_2025.

## Results

### Identifying modulators of TopBP1 condensation with an optogenetic approach

Resistance to conventional chemotherapies is the major cause of therapy failure in patients with cancer. In CRC, irinotecan resistance is often caused by overexpression of efflux pumps (42). However, additional pathways may also contribute, particularly the DNA damage response pathways that are frequently activated in various cancer types to cope with increased DNA replication stress. TopBP1 has been implicated in oxaliplatin resistance in gastric cancer (43) and is overexpressed in several cancer types (44). We first examined the number of TopBP1 condensates in the CRC cell lines HCT116 and HCT116-SN50 (50-fold more resistant to SN-38, the active metabolite of irinotecan, than the parental HCT116 cell line) (45) and observed a significantly higher number of TopBP1 foci in HCT116-SN50 cells (**Figure S1**). Moreover, our previous findings suggest that TopBP1 condensation is required to amplify ATR activity (26). Therefore, targeting TopBP1 assembly could be a promising strategy for cancer treatment. To this end, we developed an optogenetic system for screening molecules that modulate the formation of TopBP1 condensates. We used the stable Flp-In 293 T-Rex cell line in which expression of TopBP1 fused to the photoreceptor Cry2 and mCherry (optoTopBP1) is induced by doxycycline. Cry2 forms tetramers when exposed to 488 nm light and thereby nucleates the assembly of TopBP1 condensates (26). The advantage of this optogenetic approach is that TopBP1 condensates are formed on demand, within minutes, in the absence of DNA damaging agents that can induce confounding effects after prolonged treatment. After a 2-hour incubation step of optoTopBP1-expressing Flp-In 293 T-Rex cells with 10 µM of each molecule from the TargetMol library of preclinical and clinical compounds, we exposed cells to an array of blue light-emitting diodes for 5 min of light-dark cycles (4 s light followed by 10 s dark) to induce optoTopBP1 condensation (**Figure 1A**). After quantification of the number of TopBP1 foci per nucleus, we assigned a z-score to each candidate molecule based on this count. We considered molecules with a z-score lower than -2 as potential TopBP1 condensation inhibitors (red dots in **Figure 1B**). We identified >300 molecules that could be potential TopBP1 condensation inhibitors among the 4730 compounds. Then, we selected 131 compounds (based on the best z-scores) and assessed their effect on the ATR pathway activation in HCT116 cells using two complementary approaches. First, we evaluated these compounds at two low concentrations (1 and 10 µM) using a quantitative immunofluorescence assay (46) in flat-bottittiomed 384-well plates to monitor phosphorylation at S315 of the kinase effector Chk1 following incubation with SN-38 alone or with each of the 131 pre-selected compounds (**Figure 1C**). This allowed us to select the ten most effective drugs (MCB-613, OTSSP167, AZD2858, P005091, Tubercidin, Dactolisib, Physalin F, PFK158, PF562271, Torkinib) based on their ability to inhibit Chk1 phosphorylation. For the second approach, we incubated HCT116 cells with 5-Fluorouracil (5-FU) and SN-38 (FOLFIRI thereafter) to mimic *in vitro* the first-line chemotherapy regimen for patients with metastatic CRC (i.e., FOLFIRI: folic acid or leucovorin, 5-FU, and irinotecan) and each of the ten drugs and detected their interactions using a full-range dose matrix approach and SRB cytotoxicity assays. We quantified cell survival and the additive, synergistic or antagonistic effects. We observed the strongest synergistic interactions between FOLFIRI and AZD2858 (a known GSK-3β inhibitor). Consequently, we focused our study on AZD2858, which was efficient already at 1 µM (**Figure 1D**).

**Figure 1.**
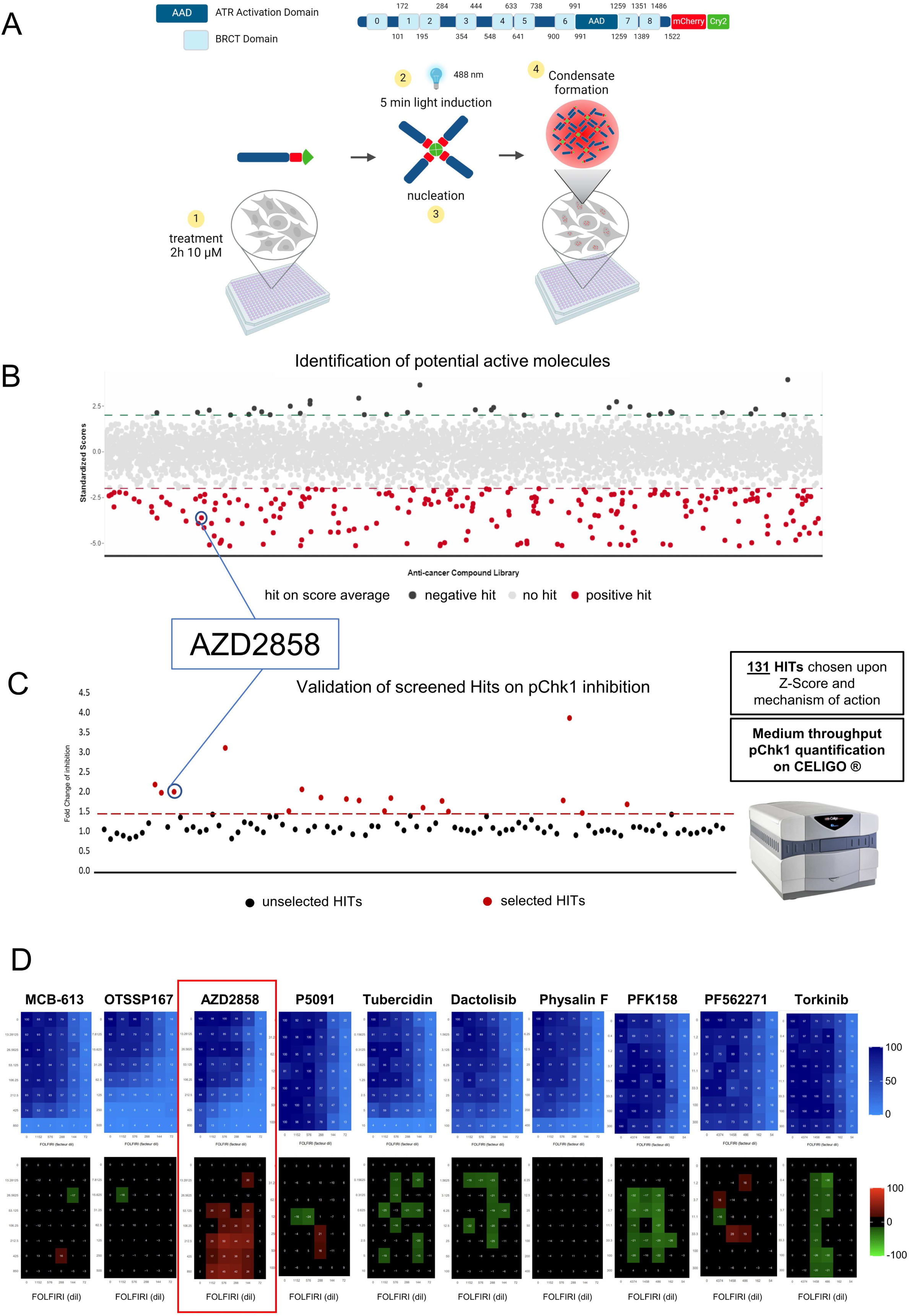
Identification of TopBP1 condensation inhibitors by high throughput screening. (**A**) Schematic description of the high-throughput screening system. Doxycycline-inducible TopBP1 was fused to mCherry and the light-sensitive cryptochrome 2 (Cry2) at its C-terminus and stably integrated in Flp-In 293 T-Rex cells. A 5-min blue light exposure (cycles of 4 s on and 10 s off) allows inducing optoTopBP1 condensation in the absence of DNA damage. Cells were grown in 384-well plates, incubated with 10 µM TargetMol molecules for 2 h before inducting optoTopBP1 condensate formation. (**B**) Graphical representation of the screening results. Drugs with a z-score lower than -2 were considered inhibitors, and drugs with a z-score higher than 2 were considered activators of TopBP1 condensate formation. The screening was performed in triplicate and each dot represents the mean z-score of a molecule. (**C**) Graphical representation of the screening confirmation in HCT116 cells. Chk1 phosphorylation at S345 was assessed using Celigo® immunofluorescence imaging after 2 h co-incubation with SN-38 and each of the 131 drugs selected from the first screening. Drugs leading to Chk1 phosphorylation (pChk1) inhibition (>1.5-fold change) were considered promising candidates and the ten molecules leading to the highest inhibition were selected for the next screening. (**D**) Viability and synergy matrices obtained after HCT116 cell incubation with increasing concentrations of FOLFIRI (5FU from 0.009 to 0.148 µM, SN38 from 0.077 to 1.235 nM) and each of the ten drugs (MCB-613 13.3 to 850 nM, OTSSP167 7.8 to 500 nM, AZD2858 13.3 to 850 nM, P005091 31.5 to 1000 nM, Tubercidin 0.15 to 10 nM, Dactolisib 1.58 to 100 nM, Physalin F 6.25 to 400 nM, PFK158 1.2 to 300 nM, PF562271 1.2 to 300 nM, Torkinib 0.4 to 300 nM) for 96 h. Cell viability was assessed with the SRB assay. Blue matrices represent cell viability. The black/red/green matrices represent additivity/synergy/antagonism, respectively. The synergy matrices were calculated with an R script (see Materials and Methods).

### AZD2858 hampers SN-38-induced endogenous TopBP1 condensate formation and ATR activation

We then evaluated AZD2858 capacity to modulate endogenous TopBP1 condensation in HCT116 cells. To this aim, we induced TopBP1 condensation by incubating cells with 300 nM SN-38 or/and 100 nM AZD2858 for 2 h. Co-incubation with AZD2858 resulted in a significant reduction in the number of endogenous TopBP1 condensates per nucleus (**Figure 2A**). Co-incubation with AZD2858 also led to a decrease in the number of 53BP1 foci (**Figure S2A**). 53BP1 forms biomolecular condensates (47) and might interact with TopBP1 (48,49). Conversely, co-incubation with AZD2858 did not modulate the number of PML nuclear bodies, which are membrane-less organelles unlike the Arsenic treatment, which is known to induce the formation of PML nuclear bodies (50) (**Figure S2B**). We then evaluated the effect of incubation of HCT116 cells with AZD2858 or/and SN-38 for 2 hours on ATR signaling (TopBP1 target), as well as on ATM (Ataxia Telangiectasia Mutated), another phosphoinositide 3-kinase-related protein kinase. Western blot analysis showed that SN-38-mediated ATR signaling induction was severely reduced by co-incubation with 100 nM AZD2858, as indicated by the decrease in Chk1 phosphorylated at S345, and RPA32 phosphorylated at S33 (**Figure 2B**). We obtained a similar result for Chk1 phosphorylated at S345 using the Celigo cytometer in cells incubated with AZD2858 and FOLFIRI (**Figure 2C**). Co-incubation of AZD2858 with SN-38 or FOLFIRI did not affect the ATM pathway, as indicated by the unchanged levels of ATM phosphorylated at S1981 and Chk2 phosphorylated at T68 (**Figure 2B and S3A)**. The level of γH2AX, a marker of DNA damage, was not affected by co-incubation with AZD2858 and SN-38 (or FOLFIRI) compared with SN-38 (or FOLFIRI) alone (**Figure 2B-C).** We obtained similar results (i.e. decrease in Chk1 phosphorylated at S345 and unchanged γH2AX levels) after co-incubation with FOLFIRI+AZD2858 for 20 hours (**Figure S3B**). Altogether, these findings suggest that AZD2858 significantly affects the ATR/Chk1 signaling pathway by inhibiting the assembly of TopBP1 condensates induced by the DNA-damaging agent SN-38.

**Figure 2.**
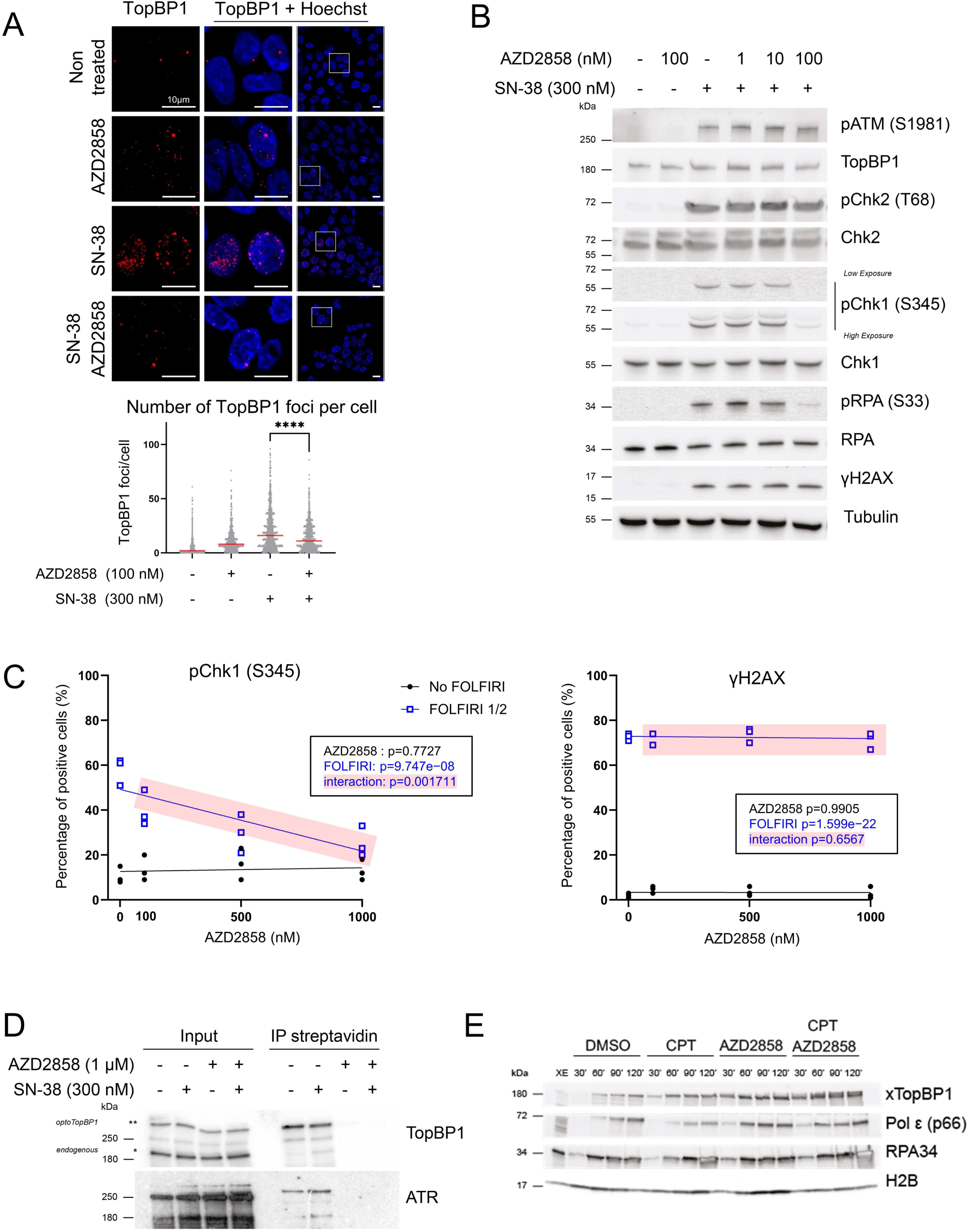
AZD2858 inhibits the formation of endogenous TopBP1 condensates and the ATR/Chk1 signaling pathway. (**A**) Representative immunofluorescence images (upper panels) of TopBP1 condensates in HCT116 cells incubated with AZD2858 (100 nM) or/and SN-38 (300 nM) for 2 h, and the corresponding quantification (lower panel). Scale bars: 10 µm. The experiment was replicated 3 times. Cell profiler was used for quantifying TopBP1 foci (>1000 cells analyzed per condition). Statistical significance was first assessed using ANOVA (p-value < 2.2e-16). The Mann-Whitney test, represented in the figure, was then used to specifically compared SN-38 and AZD2858+SN-38 conditions (****: p < 0.0001). For more details on the analyses, refer to the Materials and Methods section. (**B**) Immunoblot of the indicated proteins after incubation of HCT116 cells with AZD2858 at the indicated concentrations and/or SN-38 (300 nM) for 2 h. The experiment was replicated 3 times, and a representative replicate is shown. (**C**) Percentage of HCT116 cells positive for phosphorylated Chk1 (S345) or γH2AX expression in function of the AZD2858 concentration (0 nM, 100 nM, 500 nM, 1000 nM) in the presence (blue) or not (black) of FOLFIRI (dilution: 1/2, corresponding to 6 µM of 5-FU and 50 nM of SN-38). Cells were incubated for 2 h and positive cells were identified using Celigo® immunofluorescence imaging. The experiment was replicated 3 times, and each point represents a biological replicate. The statistical significance was determined by linear modeling interrogating the effect of each drug separately and their interaction (*p*-values given in the inset). (**D**) Immunoblotting of TopBP1 and ATR isolated with streptavidin beads from optoTopBP1-expressing cells incubated with doxycycline for 16 h to induce optoTopBP1 expression, and also with AZD2858 (1 µM) and/or SN-38 (300 nM) for the last 2 h. Biotin was added to the medium in all conditions for the last 30 min. Bands that correspond to endogenous and optoTopBP1 proteins are indicated with one and two stars (* and **), respectively. The experiment was replicated twice, and a representative result is shown. (**E**) Chromatin was extracted at the indicated time points (min) following the assembly of nuclei from sperm DNA incubated in *X. laevis* egg extracts incubated with DMSO (control), camptothecin (CPT; 55 µM) or/and AZD2858 (1 µM). Chromatin samples were analyzed by western blotting.

### AZD2858 modulates ATR/Chk1 signaling by disrupting TopBP1 assembly

As AZD2858 is a known GSK-3β inhibitor, we asked whether its effects on the ATR/Chk1 signaling pathway might be mediated by inhibiting the GSK-3β pathway. We incubated SW620 cells in which endogenous GSK-3β was depleted or not by shRNA with SN-38 or/and AZD2858 (0.1 and 1 µM) for 2 hours. SN-38-induced Chk1 phosphorylation at S345 was comparable in GSK3-β-depleted cells (shGSK3) and control cells (shLuc) (**Figure S4A**). Moreover, the SN-38 and AZD2858 combination inhibited Chk1 phosphorylation in shGSK3 and shLuc cells, indicating that GSK-3β is not involved in Chk1 phosphorylation inhibition by AZD2858. The combination did not increase significantly the level of GSK-3β phosphorylated at S9, a marker of kinase inhibition, unlike what observed with insulin, a well-known regulator of GSK-3β activity (51) (**Figure S4B**). In addition, among the molecules tested from the TargetMol library, other more effective and specific GSK-3β inhibitors, compared with AZD2858, did not show any effect on light-induced optoTopBP1 condensation (**Table S1**). These findings suggest that although AZD2858 is a GSK-3β inhibitor, its modulation of the ATR/Chk1 pathway is not due to this function. To identify the underlying mechanisms, we used a biotin proximity-labeling approach to assess TopBP1 interaction with cellular partners in the presence of AZD2858 or/and SN-38 (52). For this, we used our optoTopBP1 construct tagged at its N-terminus with TurboID, an optimized biotin ligase that allows protein biotinylation within minutes (53). We incubated Flp-In 293 T-Rex cells that stably express TurboID-tagged optoTopBP1 with AZD2858 or/and SN-38 for 2 hours and added biotin for the last 30 minutes. Isolation of biotinylated proteins using streptavidin-coated beads highlighted a drastic reduction in TopBP1 proximity with recombinant and endogenous TopBP1 and also with ATR upon incubation with AZD2858 alone or with SN-38 (**Figure 2D**). These findings suggest that AZD2858 disrupts TopBP1 self-association and its interaction with its primary activator ATR. This may contribute to the observed decrease in SN-38-induced TopBP1 foci following AZD2858 administration. Next, we determined whether TopBP1 recruitment to chromatin was altered by AZD2858 using the well-established cell-free DNA replication system derived from *X. laevis* egg extracts in which replication of demembranated sperm nuclei occurs synchronously, independently of transcription or protein synthesis. In the presence of the topoisomerase I inhibitor CPT, replication fork progression was challenged, leading to a reduction in polymerase (Pol) ε levels and an accumulation of DNA-bound TopBP1 and RPA to activate the checkpoint response (**Figure 2E**). When AZD2858 was added to the extract alone or with CPT, TopBP1 recruitment to chromatin was not inhibited. Our data demonstrate that combining CPT and AZD2858 earlier enhances the accumulation of replication-related factors (RPA, TopBP1, and Pol ε) on chromatin compared to CPT treatment alone, particularly visible at the 60-minute after starting replication (**Figure 2E**). This may be due to increased origin activation, as normally seen when the S-phase checkpoint is inhibited (54). Collectively, these observations suggest that AZD2858 acts primarily on the ATR signaling pathway through TopBP1 self-association inhibition.

### AZD2858 inhibits induction of the S-phase checkpoint by SN-38

Chk1 plays a pivotal role as a mediator of the S-phase checkpoint (55). As SN-38-induced phosphorylation of Chk1 was decreased after incubation with AZD2858, we investigated the impact of 2 hours treatments of SN-38 or/and AZD2858 on the cell cycle distribution using a BrdU/PI flow cytometry assay. BrdU incorporation was markedly reduced in SN-38-treated HCT116 cells, suggesting a slowdown of the S phase (**Figure 3A**). However, when SN-38 was combined with AZD2858, this effect was partially reduced (red arrows in **Figure 3A**).

**Figure 3.**
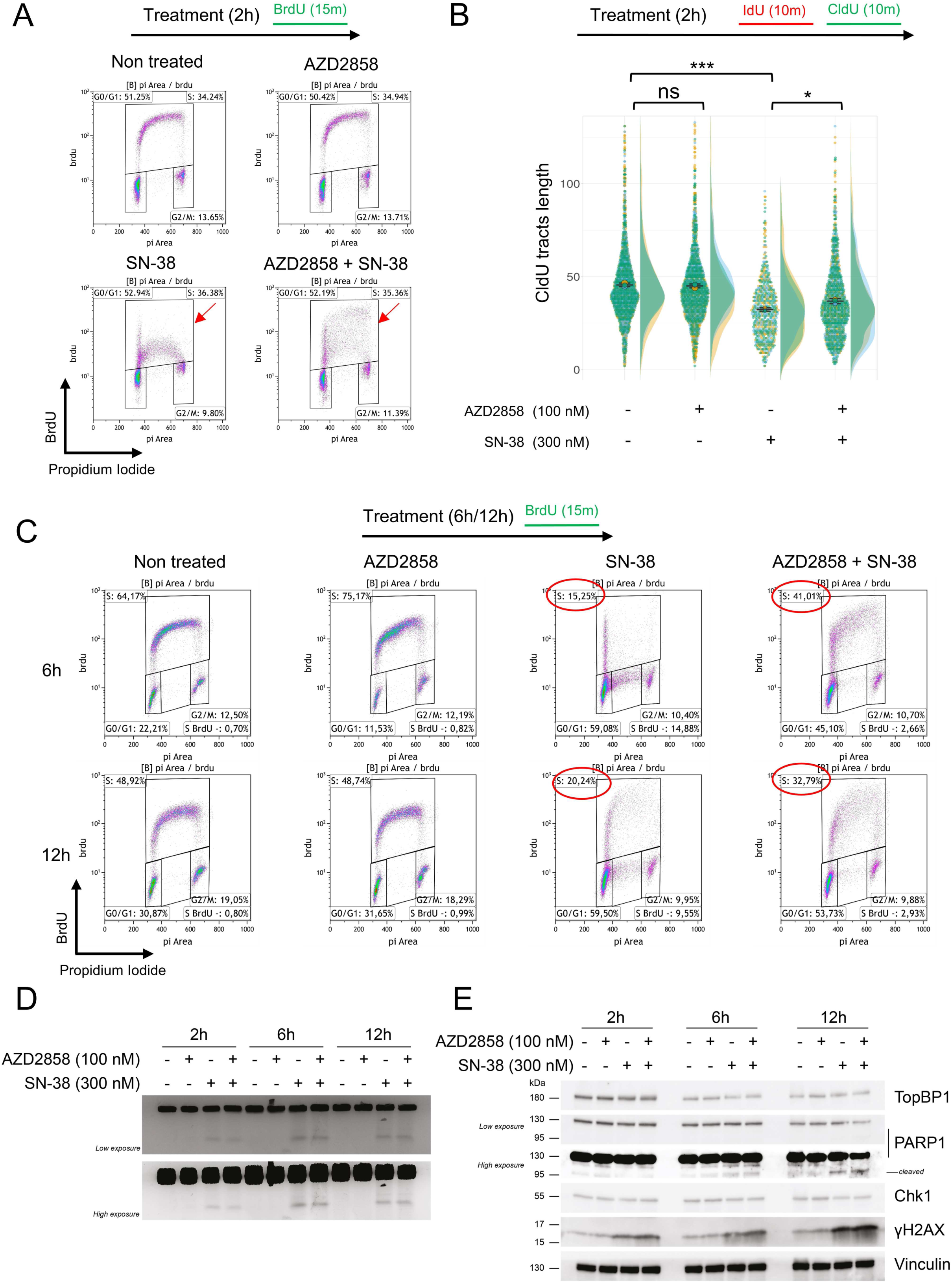
AZD258 inactivates the S-phase checkpoint in SN-38-treated cells. (**A**) Cell cycle analysis after a 2-h incubation with AZD2858 (100 nM) and/or SN-38 (300 nM); 10 µM BrdU was added in the last 15 min of the 2-h incubation. Cell debris were gated out and BrdU incorporation was plottittied against DNA content (stained with PI). Red arrows indicate BrdU incorporation. The gates for the S-phase population (BrdU-positive cells) were set broadly to prevent bias and ensure inclusion of cells with weak BrdU incorporation, particularly in the SN-38-only condition. The percentage of cells within this gate remains comparable across conditions, even though the FACS plot’s overall shape changes, reflecting a shift in BrdU incorporation distribution and highlighting a heterogeneous population with varying levels of BrdU incorporation, especially in the combination treatment. 20,000 events per condition were analyzed. The G_0_/G_1_, S and G_2_/M gates are shown. The experiment was replicated 3 times, and a representative replicate is shown. More information is available in **Table S2.** (**B**) DNA fiber analysis of replication tracts labeled with two sequential 15-min pulses of IdU and CldU added at the end of the 2-h incubation with AZD2858 (100 nM) +/-SN-38 (300 nM). The dot plots show the CldU tract length of individual replication forks. Data are pooled from n = 3 biological replicates (yellow, blue, green). Data distribution is shown as half-violin plots. Circles represent the mean of each replicate, and the error bars represent the SEM of the means of the three replicates. Statistical significance was first assessed using ANOVA (p-value < 2.2e-16). The t-test, represented in the figure, was then used to specifically compared the indicated conditions (ns: non-significant; *: p<0.05; ***: p<0.001). For more details on the analyses, refer to the Materials and Methods section. (**C**) Same as (A) but after 6 h or 12 h of incubation with AZD2858 (100 nM) and/or SN-38 (300 nM). The gate of interest (S-phase) and the corresponding cell percentages are outlined in red. More information is available in **Table S3 and Figure S5C and S5D** (**D**) PFGE analysis of DNA damage in HCT116 cells incubated with AZD2858 (100 nM) and/or SN-38 (300 nM) for the indicated times. The experiment was replicated 3 times, and a representative replicate is shown. (**E**) Immunoblot of the indicated proteins after incubation with AZD2858 (100 nM) or/and SN-38 (300 nM) for the indicated times. The experiment was replicated 3 times, and a representative replicate is shown.

To determine whether this effect depends on replication fork progression or replication origin firing, we co-labeled replication tracks with two consecutive pulses of the thymidine analogs IdU and CldU (**Figure 3B**). As expected, CldU tracks were shorter in HCT116 cells incubated with SN-38 alone, while addition of AZD2858 significantly reduced the negative impact of SN-38 on replication fork progression. We observe a heterogeneous population in the combination treatment, with some CldU tracts remaining short, similar to those in SN-38-treated cells, while others were longer, resembling those observed in untreated cells. This heterogeneity contributes to an overall increase in the mean length of CldU tracts in cells treated with the combination of AZD2858 and SN-38.

Moreover, the impact of the combination treatment becomes more pronounced over time. Indeed, there was a tendency for the SN-38-induced S-phase checkpoint to be weakened after 6 and 12 hours of co-treatment with AZD2858 and SN-38. (**Figure 3C** and **Figure S5C-D**). In the representative biological replicate shown, the percentage of BrdU incorporating cells increased from 15% (SN-38 alone) to 41% (combination) at 6 hours and from 21% (SN-38 alone) to 34% (combination) at 12 hours (**Figure 3C** and **Figure S5C-D**). These results indicate that the combination of AZD2858 and SN-38 affects replication fork progression and disrupts the cell cycle homeostasis, likely by inhibiting the ATR/Chk1 signaling pathway.

To further investigate the impact of AZD2858 on cell cycle progression, we performed a pulse-chase experiment in HCT116 cells. Cells were pulse-labeled with BrdU for 15 minutes, washed and then incubated with SN-38 and/or AZD2858 for 6 and 12 hours. In the combination treatment, at 12 hours, we observed an increase in the proportion of cells transitioning into S phase, which were initially in G1 (BrdU-negative) and progressed to a mid-S DNA content state (between 2N and 4N) (green squares) (**Figure S5A** and **S5B**). This observation aligns with previous findings indicating that the S-phase checkpoint is partially alleviated in the combination treatment compared to SN-38 alone. Of note, in untreated cells and those treated with AZD2858 alone, the S-phase gate represents a new round of DNA synthesis, as indicated by the progression from the new G1 phase at 6 hours to the initiation of a new S phase at 12 hours. However, in cells treated with SN-38 or the combination of SN-38 and AZD2858, the same S-phase population observed at 6 hours persists at 12 hours, reflecting a marked slowing of cell cycle progression.

Finally, the combined treatment led to a progressive accumulation of BrdU-positive cells with 4N DNA content, corresponding to cells that completed S phase (BrdU-positive) and progressed into G2 phase (4N DNA content) (red squares) (**Figure S5A** and **S5B**). This effect was more pronounced after 12 hours of treatment, with the proportion of BrdU-positive cells in G2 phase increasing from 4% in SN-38-treated cells to 11% in the combination treatment, as shown in the representative experiment in **Figure S5B**. While this trend was consistently observed across all experiments, the increase was modest and did not reach statistical significance (**Figure S5F**). This suggests that the combination of SN-38 and AZD2858 attittienuates the S-phase checkpoint, thereby activating a compensatory post-replicative cell cycle checkpoint that arrests the cell cycle before mitosis.

Abrogation of the replication checkpoint allows cells to enter and progress into the S phase with unrepaired DNA damage. We evaluated the level of DNA double-strand breaks (DSBs) using PFGE and monitored γH2AX and cleaved PARP1 expression by western blotting in cells exposed to SN-38 and AZD2838 at various time points. DSBs remained elevated after the combined treatment (**Figure 3D**). Additionally, γH2AX and cleaved PARP1 levels increased at later time points of incubation (**Figure 3E**). Altogether, these findings indicate that the AZD2858 and SN-38 combination disrupts TopBP1 assembly, most likely by inhibiting ATR kinase activity amplification, a critical process for inducing the S-phase checkpoint and regulating DNA damage repair.

### AZD2858 treatment synergizes with conventional chemotherapy drugs

In the absence of an effective S-phase checkpoint, DNA damage can lead to apoptosis and cell death. To assess this, we stained HCT116 cells with PI and quantified the sub-G1 population to measure cell death. After 48-hour incubation with sub-lethal doses of SN-38 and AZD2858, the sub-G1 cell population was increased by 40% compared with untreated cells, indicative of apoptotic cells (**Figure 4A**) and consistent with the early increase of cleaved PARP1 (**Figure 3E**). We obtained the same results in cells incubated with AZD2858 and FOLFIRI. Specifically, in the representative experiment shown in **Figure 4A**, the sub-G1 population increased from 19% after incubation with SN-38 alone to 61% after incubation with SN-38+AZD2858, and from 27% after incubation with FOLFIRI alone to 71% after incubation with FOLFIRI+AZD2858 (**Figure 4B**). Additionally, the combined treatments (AZD2858 with SN-38 or FOLFIRI) led to an increase of cleaved PARP1 and cleaved caspase-3 (western blotting, **Figure 4C**), and to the accumulation of DNA DSBs (PFGE, **Figure 4D**). Then, we performed an SRB assay in HCT116 cells incubated with the AZD2858 and SN-38 combination, as previously described, and visualized the viability (blue) and synergy (black/red) matrices. The AZD2858 and SN-38 combination, like the AZD2858 and FOLFIRI combination (**Figure 1D**), synergistically inhibited cell survival (**Figure 4E**).

**Figure 4.**
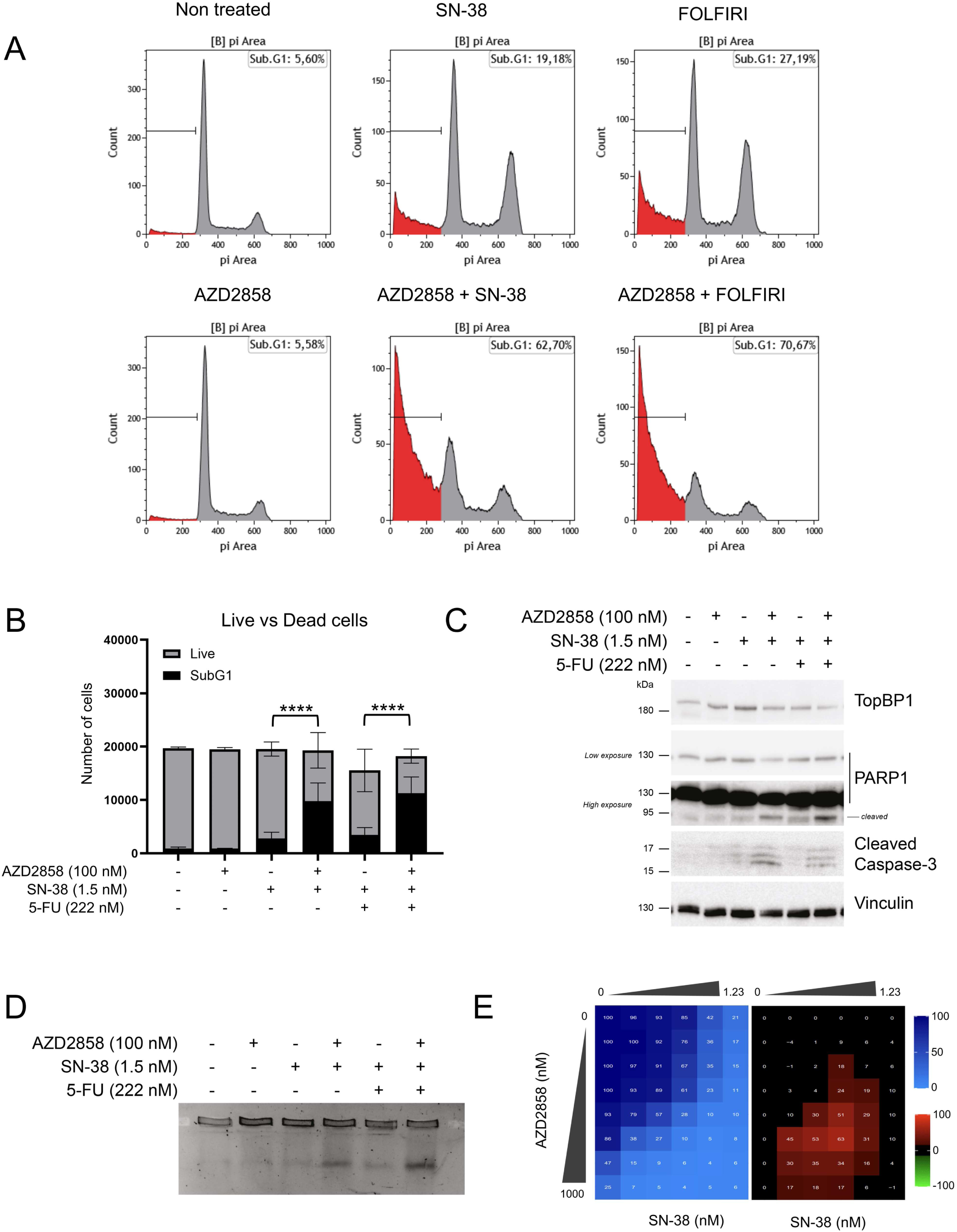
AZD2858 synergizes with SN-38 and FOLFIRI by inducing DNA damage and cell apoptosis. (**A**) Flow cytometry analysis to quantify sub-G1 HCT116 cells after 48-h incubation with sub-optimal doses of AZD2858 (100 nM) alone or with SN-38 (1.5 nM) or FOLFIRI (SN-38 1.5 nM; 5-FU 222 nM). The x-axis shows the DNA content (PI staining) and the y-axis the cell count. The presence of sub-G1 cells gated under the G1 peak suggests DNA fragmentation, a characteristic feature of apoptotic cell death. The experiment was replicated 3 times, and a representative replicate is shown. (**B**) Graph showing the number of sub-G1 cells (considered as dead cells) versus the number of live cells, according to the indicated treatments (from A). Error bars represent 3 individual biological replicates. The statistical significance was determined by linear regression analysis, more information is available in **Table S4** (****: p<0.0001). (**C**) Immunoblot of the indicated proteins in HCT116 cells after 48-h incubation as described in A. (**D**) PFGE analysis of DNA damage in HCT116 cells incubated for 48 h as described in A. (**E**) HCT116 cells were incubated with increasing concentrations of SN-38 (0 to 1.23 nM) and AZD2858 (0 to 1000 nM) for 96 h (2D culture system). Cell viability was assessed with the SRB assay. The synergy matrix was calculated as described in Materials and Methods. The experiment was replicated 3 times, and a representative replicate is shown.

Next, we assessed whether the combination affected cell death or proliferation. First, we determined whether AZD2858, SN−38, and FOLFIRI alone induced cytotoxic (cell death) or cytostatic (inhibition of cell proliferation) effects by PI/Hoechst labeling of cells incubated with increasing concentrations of each drug. The three drugs displayed mainly cytostatic effects (**Figure S6A**), as indicated by the absence of cell death (lower panels) even at doses that decreased cell proliferation (upper panels). Next, we stained with PI/Hoechst and performed SRB assays in HCT116 cells, CT−26 cells (murine CRC cell line) and CCD 841 CoN (untransformed colorectal cells) incubated with increasing concentrations of AZD2858 and FOLFIRI (matrix analysis). We found that the combination was cytostatic at low drug doses and became cytotoxic at higher doses (**Figure S6B and S6C**), especially in areas where we detected synergy (**Figure 5A and S6D).** However, no synergistic effect was observed on CCD 841 CoN cells with the AZD2858 and FOLFIRI combination (**Figure S7**). These results indicate that when the AZD2858+FOLFIRI combination shows synergistic effects, this interaction is cytotoxic. We confirmed the synergy of the FOLFIRI+AZD2858 combination in three additional human CRC cell lines: HT29, SW620 and SW480 (**Figure 5C**). We also performed a matrix analysis in two SN−38−resistant cell lines: HCT116 SN−6 and HCT116 SN−50 (56) (**Figure 5D and 5F**). The FOLFIRI+AZD2858 combination was synergistic in both SN−38−resistant cell lines, indicating that this combination may alleviate drug resistance in colorectal cancer.

**Figure 5.**
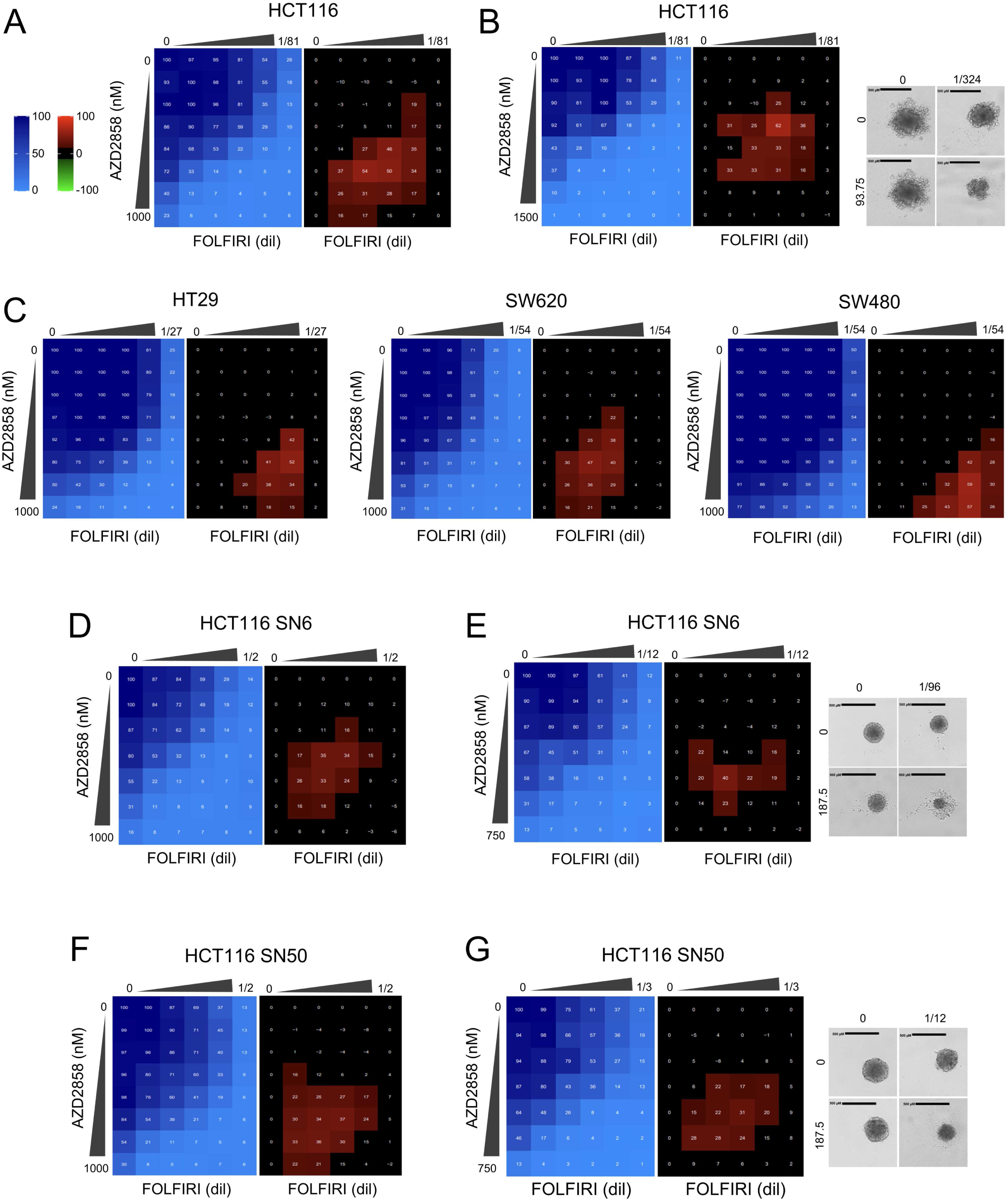
AZD2858 synergizes with FOLFIRI on a panel of CRC cell lines, including HCT116 resistant to SN-38. (**A**) HCT116 cells were incubated with increasing concentrations of FOLFIRI (5-FU from 0.009 to 0.148 µM and SN-38 from 0.077 to 1.235 nM) and AZD2858 (from 15.6 to 1000 nM). Cell viability was assessed with the SRB assay in 2D cultures to generate the viability matrix (blue). The synergy matrices (black and red) were calculated as described in Materials and Methods. (**B**) HCT116 cells were cultured in 3D to form spheroids and incubated with increasing concentrations of FOLFIRI (5-FU from 0.009 to 0.148 µM and SN-38 from 0.077 to 1.235 nM) and AZD2858 (from 23.4 to 1500 nM). Cell viability was assessed with CellTiter-Glo to obtain the viability matrix (blue). The synergy matrices (black and red) were calculated as described in Materials and Methods. Representative brighvield images of spheroids are shown: untreated and after incubation with the drug concentrations giving the highest synergy score. (**C**) HT29, SW620 and SW480 CRC cells were incubated (2D culture) with increasing concentrations of FOLFIRI (HT29 cells: 5-FU from 0.03 to 1 µM and SN-38 from 0.07 to 2.35 nM; SW620 and SW480 cells: 5-FU: from 0.13 to 4.4 µM, SN-38: from 0.007 to 0.25 nM) and AZD2858 (from 15.6 to 1000 nM). (**D**) SN-38-resistant HCT116 SN-6 (six times more resistant to SN-38 than wild-type HCT116, 2D culture) were incubated with increasing concentrations of FOLFIRI (5-FU from 0.46 to 14.85 µM and SN-38 from 0.1 to 3.375 nM) and AZD2858 (from 15.6 to 1000 nM). (**E**) Same as in B but with SN-38-resistant HCT116 SN-6 cells. Drug concentrations were as follows: FOLFIRI (5-FU from 0.083 to 1.333 µM and SN-38 from 0.694 to 11.1 nM) and AZD2858 (from 46.8 to 750 nM). (**F**) Same as (D) but with SN-38-resistant HCT116 SN-50 cells (fifty times more resistant to SN-38 than wild-type HCT116). (**G**) Same as in B but with SN-38-resistant HCT116 SN-50 cells. Drug concentrations were as follows: FOLFIRI (5-FU from 0.75 to 12 µM and SN-38 from 6.25 to 100 nM) and AZD2858 (from 46.8 to 750 nM). Experiments were replicated 3 times, and representative replicates are shown.

Lastly, we tested the FOLFIRI+AZD2858 combination (matrix analysis) in 3D cell cultures that are physiologically closer to tumor growth compared with 2D cultures and allow mimicking the complexity of solid tumors from a structural, biochemical and biophysical point of view. We prepared spheroids of parental HCT116 cells, SN-38-resistant HCT116 SN-6 and HCT116 SN-50 cells (**Figure 5B,E,G)**, and murine CT-26 cells (**Figure S6E)**. We again observed that the combination was synergistic in all tested cell lines, demonstrating remarkable efficacy with nearly complete elimination of spheroids.

## Discussion

TopBP1 assembly is required for the activation of the ATR signaling pathway, a critical step in the DNA damage response and repair. We exploited our ability to control TopBP1 condensation and ATR activation using an optogenetic approach to develop a high-throughput screening procedure for identifying modulators of TopBP1 condensation. As proof of concept, we screened a TargetMol library of 4730 preclinical and clinical compounds. Through this approach, we found that AZD2858, a known GSK3-β inhibitor, is also a potential inhibitor of TopBP1 condensation. Using a complementary approach based on the irinotecan active metabolite SN-38 to induce DNA damage in CRC cells, we confirmed that AZD2858 disrupts endogenous TopBP1 condensates, thus affecting the ATR signaling pathway.

AZD2858 inhibits GSK-3β activity and activates the canonical Wnt/β-catenin signaling cascade (57). In addition, AZD2858 has cytotoxic effects in glioma cell lines through disruption of centrosome function and mitotic failure (58), and affects glioma proliferation and survival (57). Our findings show that AZD2858, at nanomolar concentrations, alters TopBP1 condensation without influencing GSK-3β activity and does not require the presence of GSK-3β to prevent Chk1 activation by SN-38. These observations clearly rule out the involvement of the GSK-3β pathway in TopBP1 condensate formation inhibition by AZD2858.

TopBP1 is overexpressed in several cancer types, including breast cancer (59) and advanced-stage CRC as well as radiotherapy-resistant lung cancer cells and oxaliplatin-resistant gastric cancer (43). This overexpression is associated with high-grade tumors and poor prognosis. Notably, inhibition of TopBP1 expression increases the radiosensitivity of lung cancer and brain metastases (60) and also DNA damage in oxaliplatin-resistant gastric cancer cells (43). To date, no study assessed or visualized TopBP1 condensate levels in tumors, or investigated whether their presence is linked to poor prognosis. Here, we found that TopBP1 condensates are increased in SN-38 resistant CRC cells. Many attittiempts have been made to target TopBP1 activity. Weei-Chin Lin9s team was the first to perform two large scale molecular docking screens targeting the BRCT7/8 domains of TopBP1. They identified calcein AM and 5D4 as compounds that block TopBP1 oligomerization and demonstrated their anti-cancer activity *in vivo* (61,62). Interestingly, 5D4 also disrupts TopBP1 interaction with E2F1, mutant p53, CIP2A, and MIZ1, leading to E2F1-mediated apoptosis, suppression of mutant p53 and repression of MYC activity (61).

We demonstrated that AZD2858 disrupts the interaction between TopBP1 and its canonical partner ATR and also its self-interaction. Additionally, in cell-free *X. laevis* egg extracts, AZD2858, alone or in combination with the topoisomerase I inhibitor CPT, enhanced the DNA binding occupancy of both TopBP1 and RPA32. These findings, combined with the cell cycle and DNA fiber results, suggest that AZD2858 abrogates the intra-S phase and DNA elongation checkpoints induced by SN-38, leading to unscheduled DNA synthesis. In addition, incubation with SN-38+AZD2858 enhanced DNA damage and apoptosis, as indicated by (i) the increased γH2AX levels, (ii) the persistence of DSBs (PFGE assay), (iii) the expression of cleaved PARP1 and caspase-3 (apoptotic markers), and (iv) the accumulation of sub-G1 cells. Incubation with the AZD2858+SN-38 combination mimicked the effects observed with ATR inhibitors when combined with topoisomerase I inhibitors, such as CPT, topotecan and SN-38 (62,63). For instance, the ATR inhibitors VE-821 and VE-822, sensitize cancer cells to CPT derivatives by alleviating the intra-S-phase checkpoint (62,63), and induce apoptosis mediated by caspase-3 when combined with SN-38 because cells experience extensive DNA damage due to failure of the replication checkpoint (63). Importantly, the e昀케cacy of several ATR inhibitors is currently investigated in preclinical studies or in phase I/II clinical trials as monotherapy or in combination with chemotherapy-induced replication stress (64). We also showed a synergistic effect of the AZD2858+FOLFIRI combination, including in SN-38-resistant HCT116 cell lines, due to the increased apoptosis and accumulation of DNA damage. Given the similarity of mechanism between AZD2858 and ATR inhibitors, when combined with chemotherapy (SN-38 or CPT), we hypothesize that AZD2858 could prevent the development of resistance because its mechanism of action is not to inhibit the ATR kinase but to prevent the interaction of TopBP1 with ATR. FOLFIRI is used as first-line chemotherapy regimen for patients with metastatic CRC and gastric/gastroesophageal cancer (65).

In the last decade, a major advance has been made in our understanding of the mechanisms and functions of various cytoplasmic and nuclear membrane-less organelles within cells, revolutionizing thefield of cell biology. Recent studies highlighted the importance of compartmentalizing cellular components into mesoscale structures, known as biomolecular condensates, implicated in new physiological functions and in triggering the activation of specific signaling pathways. Many research projects have incorporated condensates into the understanding of cancer pathogenesis, leading to the development of new therapeutic strategies to target cancer-associated condensates (66,67). We recently found that TopBP1 condensation acts as a molecular switch to amplify ATR activity (26), a critical pathway to tolerate the intrinsically high levels of lesions that block replication fork progression in cancer cells. Targeting TopBP1 assembly, rather than its degradation, offers an interesting strategy to selectively inhibit its role in the ATR/Chk1 signaling pathway activation, while preserving its other essential functions in replication and transcription that do not rely on its assembly.

Overall, this study brought the first insights into the rational of targeting the condensation of TopBP1, a multifunctional protein that forms biomolecular condensates in response to DNA damage to inhibit the ATR/Chk1 signaling pathway and to potentiate conventional therapies.

## Acknowledgement

In memory of Angelos Constantinou whose significant contribution to the field of DNA damage response will remain an enduring source of inspiration for our research. We acknowledge the national infrastructure France-BioImaging supported by the French National Research Agency (ANR-10-INBS-04). This work was supported by the French Institut National du Cancer INCa (PLBIO 2021), by the French Agence Nationale de la Recherche ANR (AAPG2021), and by the Fondation MSD AVENIR. We acknowledge the imaging facility MRI, member of the France-BioImaging national infrastructure supported by the French National Research Agency (ANR-10-INBS-04, <Investments for the future=).

## Author Contributions

Conceptualization: L.M, C.G, J.B, A.C; Methodology: L.M, N.V, A.A, T.E, A.Az, L.A.M, V.G, C.G, J.B; Formal analyses: L.M, N.V, A.A, T.E, A.Az, S.F, H.S, L.A.M, V.G; Investigation: L.M, A.A, N.V, C.G, J.B, A.C; Writing original draft: L.M, A.A, C.G, J.B, A.C; Writing – review and editing: L.M, A.A, T.E, C.G, J.B, A.C; Supervision: N.V, C.G, J.B, A.C; Project Administration: C.G, J.B, A.C; Funding acquisition N.V, C. G, A.C.

## Inclusion and diversity

We support inclusive, diverse, and equitable conduct of research.

## Supplementary legends

**Figure S1.**
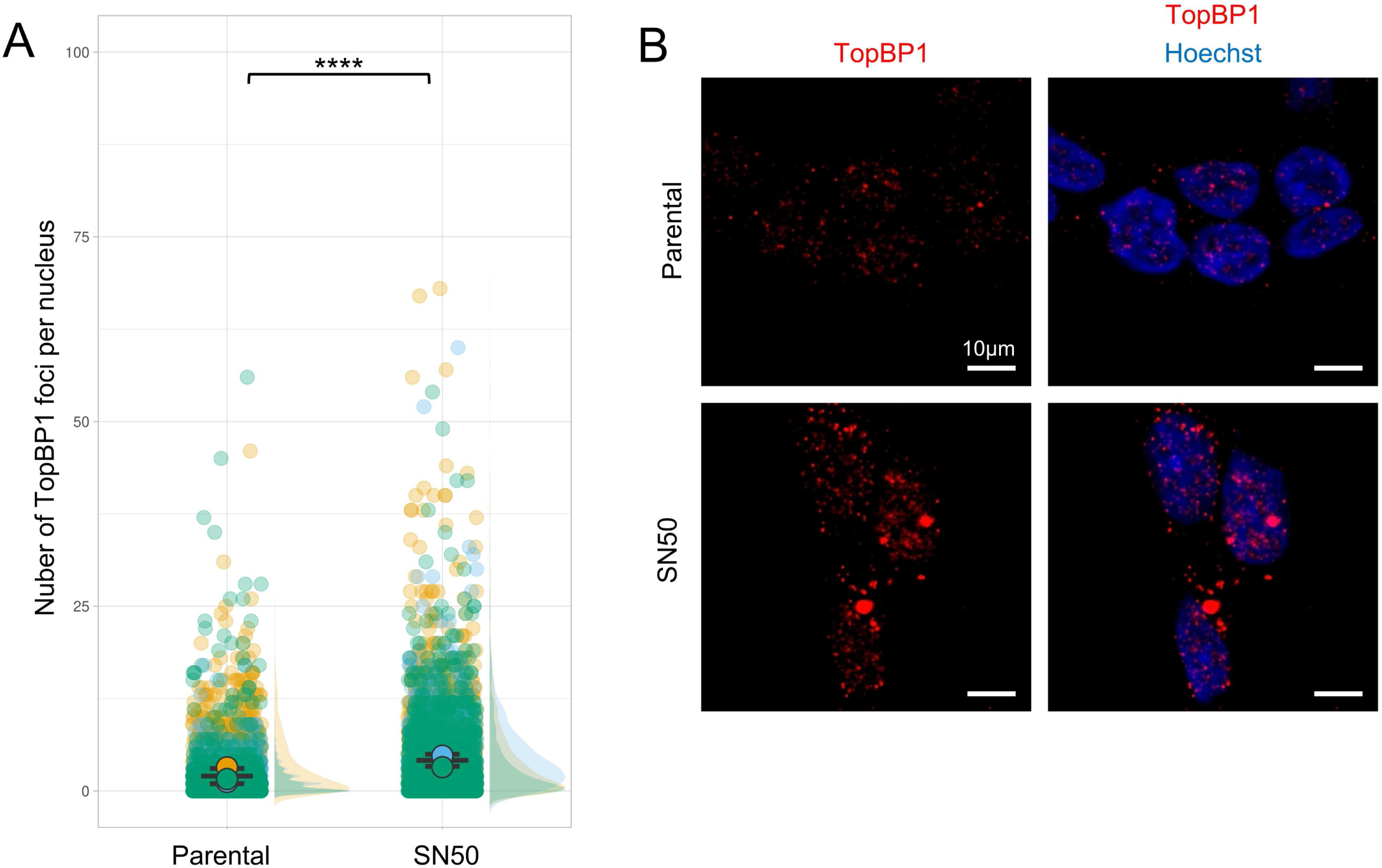
TopBP1 forms more foci in SN-38-resistant HCT116 cells. (**A**) Number of TopBP1 foci per cell in parental and SN-38-resistant HCT116 cells. Data were pooled from n = 3 biological replicates (yellow, blue, green). Data distribution is shown as a half-violin plots on the right. Circles represent the mean of each replicate, and the error bars represent the SEM of the means of the three replicates. Significance of the differences was tested using an ANOVA on a generalized linear model interrogating both the effect of cell line genotype and replicate identity (modeling TopBP1 foci number by a Poisson distribution). Foci numbers are significantly different between the two cell lines (p-value<2.2e-16) and across the 3 replicates of the experiment (p-value<2.2e-16). (**B**) Representative images of one replicate.

**Figure S2.**
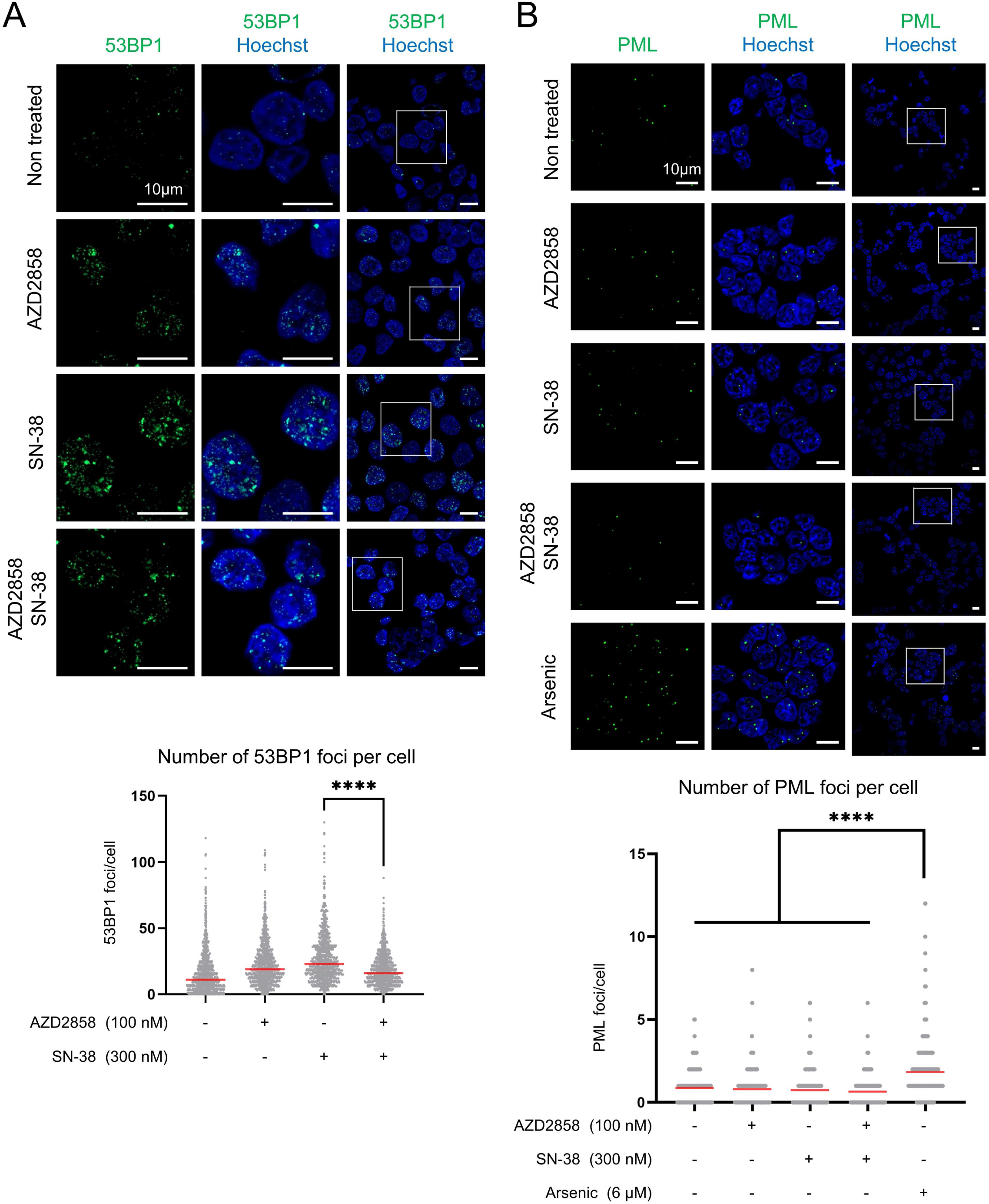
Impact of the AZD2858 and SN-38 combination on two nuclear biomolecular condensates. (A) The effect of AZD2858 (100 nM) or/and SN-38 (300 nM) on 53BP1 and (B) PML condensates were tested after 2 h of incubation. Arsenic (6 µM, 2 h) was used as a positive control for PML nuclear bodies9 induction. Scale bars: 10 µm; ****: p<0.0001 (Mann-Whitney test for 53BP1 foci, Kruskal-Wallis for multiple comparison for PML foci).

**Figure S3.**
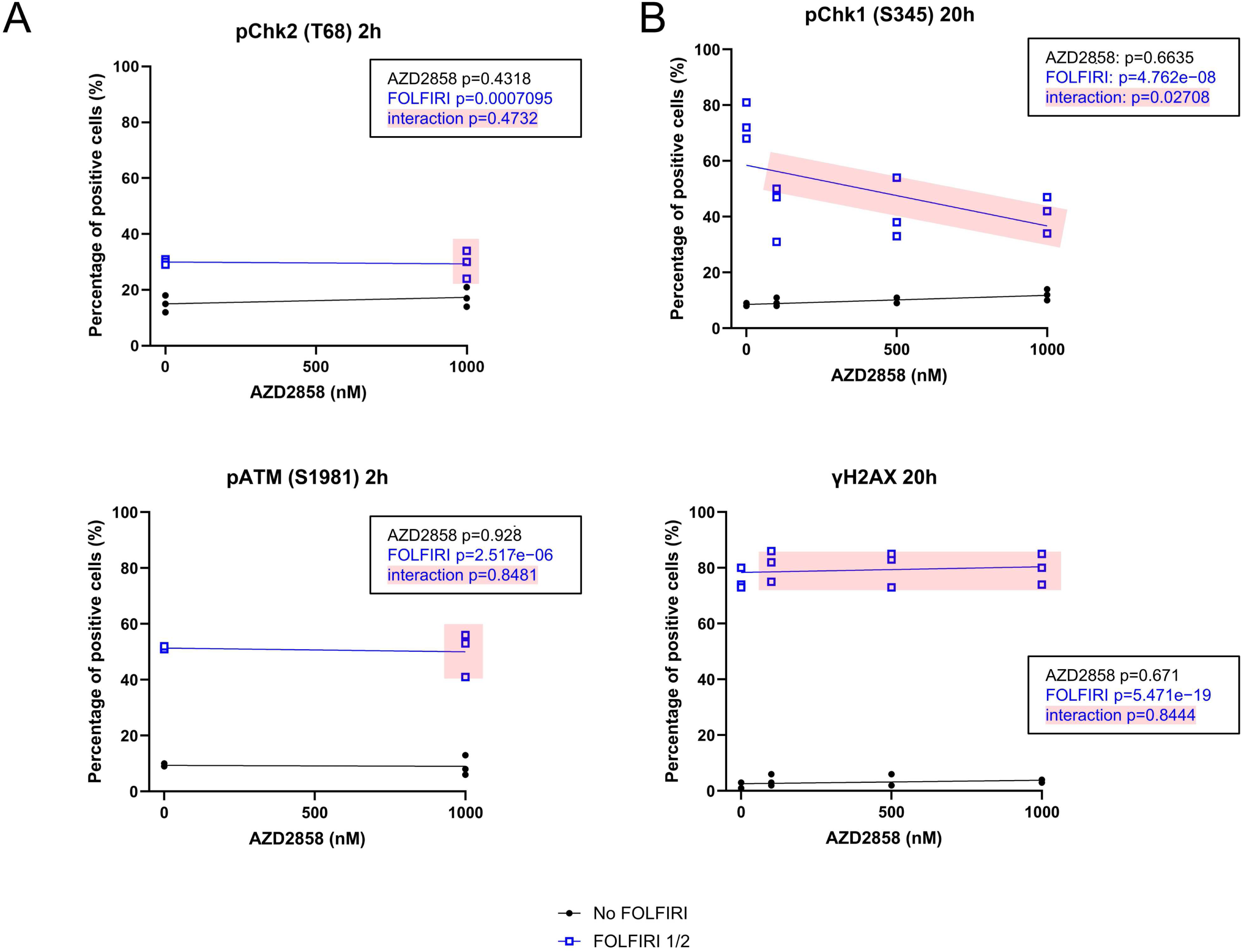
Effects of AZD2858 on components of the DNA damage response. (**A**) Quantification of the percentage of phosphorylated Chk2 (pChk2) (T68), and pATM (S1981)-positive cells in HCT116 CRC cells incubated with FOLFIRI (5-FU: 6 µM and SN-38: 50 nM) and increasing con-centrations of AZD2858 (from 250 to 1000 nM) for 2 h and (**B**) of pChk1 and γH2AX-positive cells after incubation for 20 h. Data were obtained by immunofluorescence analysis using the <Expression analysis= application of the Celigo Imaging Cytometer (Nexcelom). The combination of FOLFIRI and AZD2858 is shaded in pink. The experiment was replicated 3 times, and each point represents a biological replicate. The statistical significance was determined by linear modeling analysis interrogating the effect of each drug separately, and of their interaction (p-values given in the inset).

**Figure S4.**
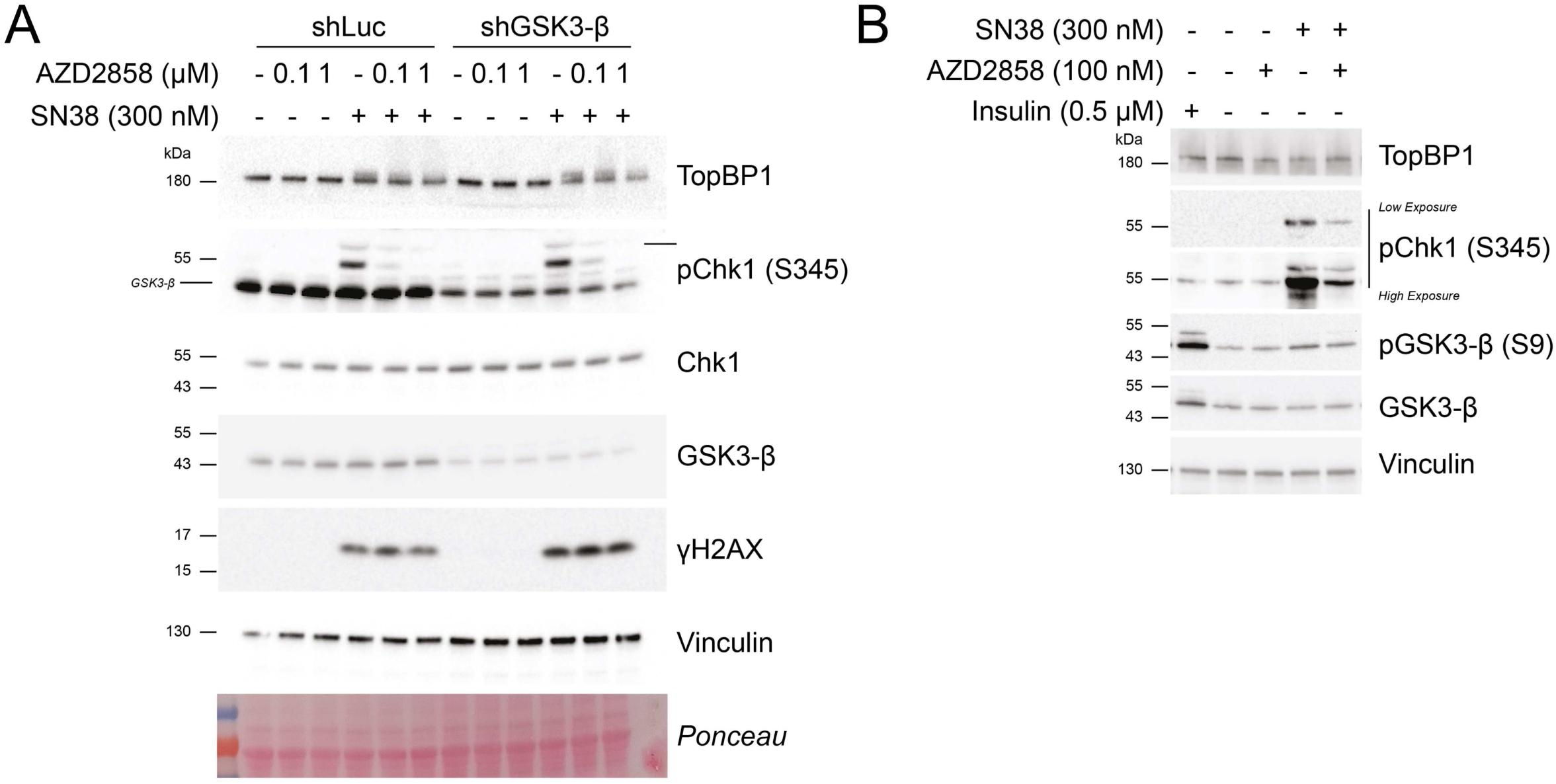
AZD2858 effect on ATR signaling is not related to GSK-3β activity. (**A**) Immunoblot of the indicated proteins after incubation of SW620 cells that express shLuc or shGSK-3β with AZD2858 (100 nM) or/and SN-38 (300 nM) for 2 h. (**B**) Immunoblot of indicated proteins after incubation of HCT116 cells with AZD2858 (100 nM) or/and SN-38 (300 nM), or insulin (0.5 µM) (positive control of GSK-3β inhibition).

**Figure S5.**
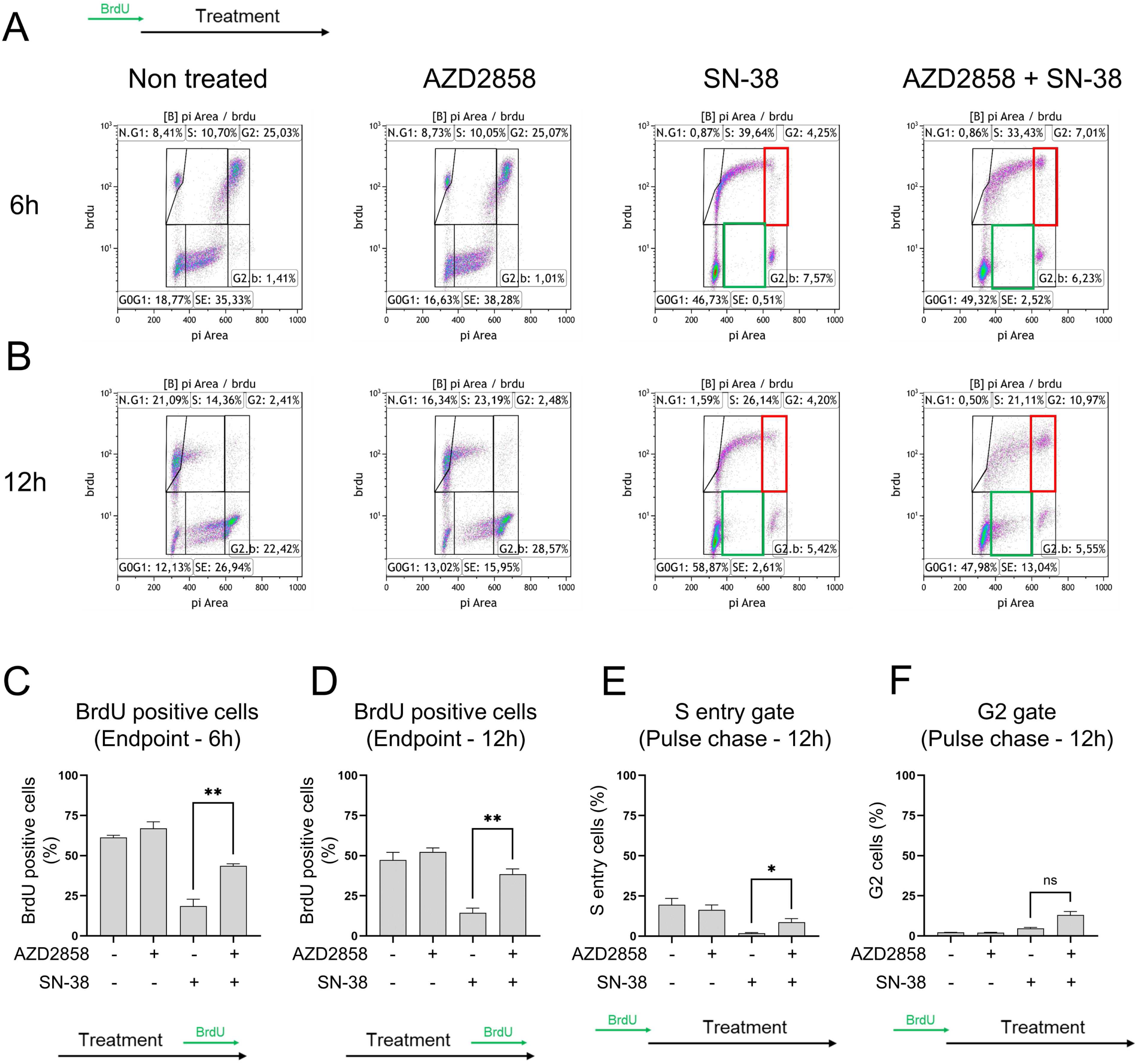
Cell cycle profiling of HCT116 cells incubated with AZD2858 or/and SN-38. (**A**) Cells were pulsed with BrdU for 15 min before the treatments, then chased without BrdU for the 6 hours of treatment, thereby named Pulse chase experiment (Non treated, AZD2858 100nM, SN-38 300nM, combination of AZD2858 and SN-38). Cellular debris were gated out and the level of BrdU incorporation was plottittied against the DNA content. 20,000 events per condition were analyzed. BrdU-positive cells include “MP” (mitotic passage, cells that were in S phase during the pulse, completed mitosis, and returned to a 2N DNA content), “S” (cells in S phase during the pulse and still in S phase at the endpoint), and “G2” (cells in S phase during the pulse that progressed to 4N DNA). Among the BrdU-negative cells, G0/G1 cells had a 2N DNA content after the chase. BrdU-negative cells with >2N DNA content entered S-phase after the BrdU pulse and were gated <SE= for S-phase entry. “G2-blocked” 4N cells were already in G2/M during the BrdU pulse and were still blocked at the endpoint. (**B**) Same as in A) but after 12 h of treatment. The gate of interest, S entry and G2 phase, are outlined in green and red. The experiment was replicated 3 times, and a representative replicate is shown. (**C**) Graphical representation of the percentage of BrdU-positive cells after 6 hours of treatment from three independent biological replicates (”Endpoint” refers to the protocol when BrdU is added at the end of the treatment, as indicated in the schematic illustration and corresponding to the representative replicate shown in Figure 3C). (**D**) Same as C) but after 12 h of treatment. (**E**) Graphical representation of three independent biological replicates of the percentage of cells in the S phase entry gate in Pulse chase experiment at 12 h of treatment (corresponding to **green squares in Figure S5B**). (**F**) Same as E but for the G2 gate (corresponding to red squares in **Figure S5B**). Statistical significance was first assessed using ANOVA (cf Materials and Mathod for detailed analysis). The t-test, represented in thefigure, was then used to specifically compared the indicated conditions (ns: non-significant; *: p<0.05; **: p<0.01 (t-test)).

**Figure S6:**
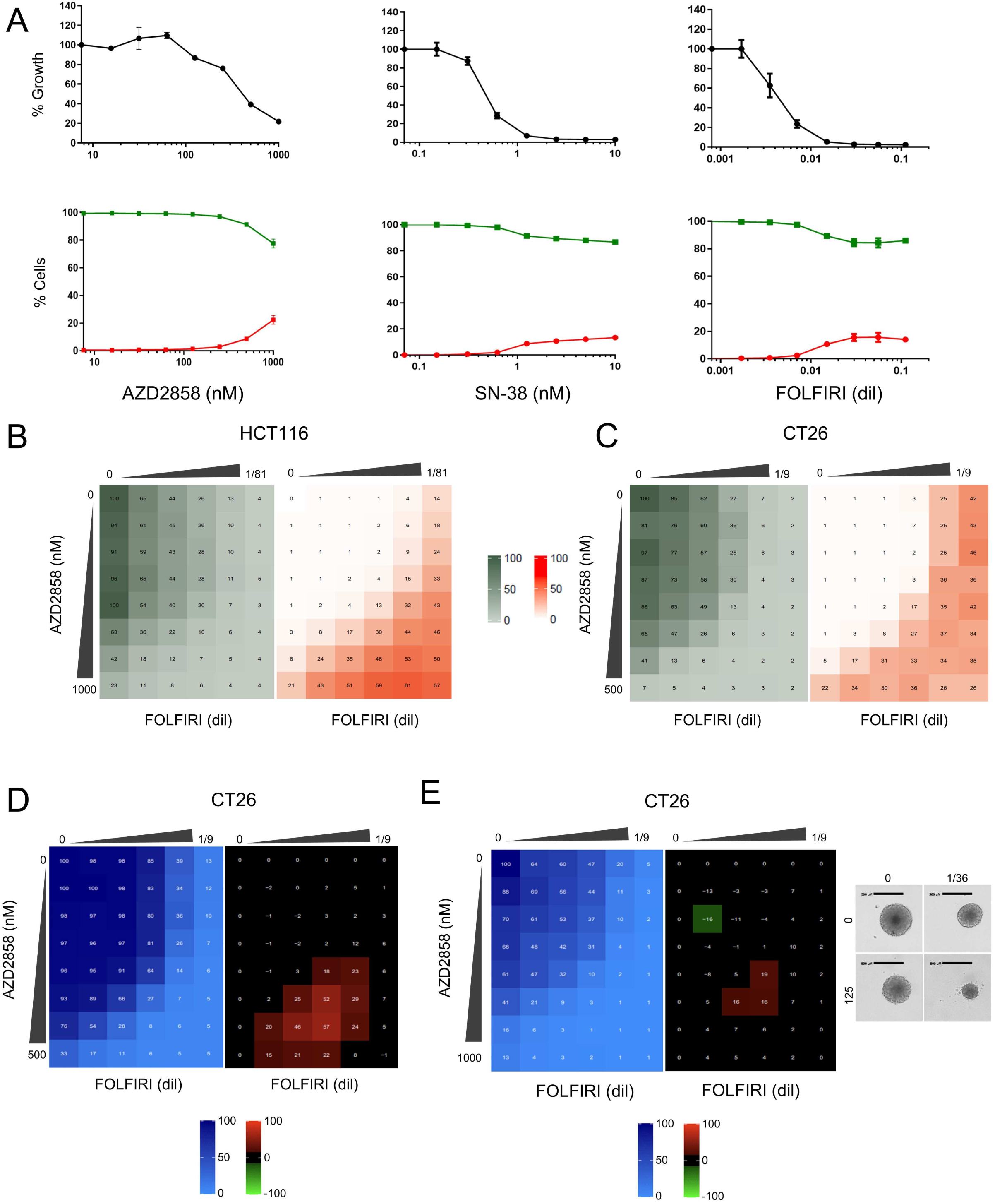
The synergistic effect of the AZD2858 + FOLFIRI combination is due to cytotoxicity. (**A**) HCT116 cells were incubated with increasing concentrations of each drug for 72 h. Then, dual staining with Hoechst/IP was performed to visualize dead cells (red) and live cells (green), respectively, in each well. Results were quantified using the dedicated Celigo® software and shown as percentages of dead cells and live cells relative to untreated controls. (**B**) HCT116 cells were incubated with increasing concentrations of FOLFIRI (5-FU from 0.009 to 0.148 µM and SN-38 from 0.077 to 1.235 nM) and AZD2858 (from 15.6 to 1000 nM) for 96 h. Then, dual staining with PI and Hoechst was performed to visualize **dead cells** (red) and **live cells** (khaki), respectively, in each well. Results were quantified using the dedicated Celigo® software: the khaki matrix represents cell survival and the orange matrix cell death, relative to untreated controls. (**C**) CT−26 cells were incubated with increasing concentrations of FOLFIRI (5−FU from 0.083 to 1.333 μM and SN−38 from 0.694 to 11.11 nM) and AZD2858 (from 7.8 to 500 nM) for 96 h. Then, dual staining with PI and Hoechst was performed to visualize dead cells (red) and live cells (khaki), respectively, in each well. Results were quantified using the dedicated Celigo® software; the khaki matrix represents cell survival and the orange matric cell death relative to un− treated controls. (**D**) CT−26 cells were incubated with increasing concentrations of FOLFIRI (5−FU from 0.083 to 1.333 μM and SN−38 from 0.694 to 11.11 nM) and AZD2858 (from 7.8 to 500 nM). Cell viability was assessed with the SRB assay in 2D cultures to generate the **viability matrix** (blue). The **synergy matrices** (black, red and green) were calculated as described in Materials and Methods. (**E**) CT−26 murine cell line cells were cultured in 3D. Drug concentrations were as follows: FOLFIRI (5−FU from 0.083 to 1.333 μM and SN−38 from 0.694 to 11.1 nM) and AZD2858 (from 15.6 to 1000 nM). Cell via− bility was assessed with CellTiter−Glo to obtain the **viability matrix** (blue). The **synergy matrices** (black and red) were calculated as described in Materials and Methods. Representative brightfield images of spheroids are shown: untreated and after incubation with the drug concentrations giving the highest synergy score. Experiments were replicated 3 times, and representative replicates are shown.

**Figure S7:**
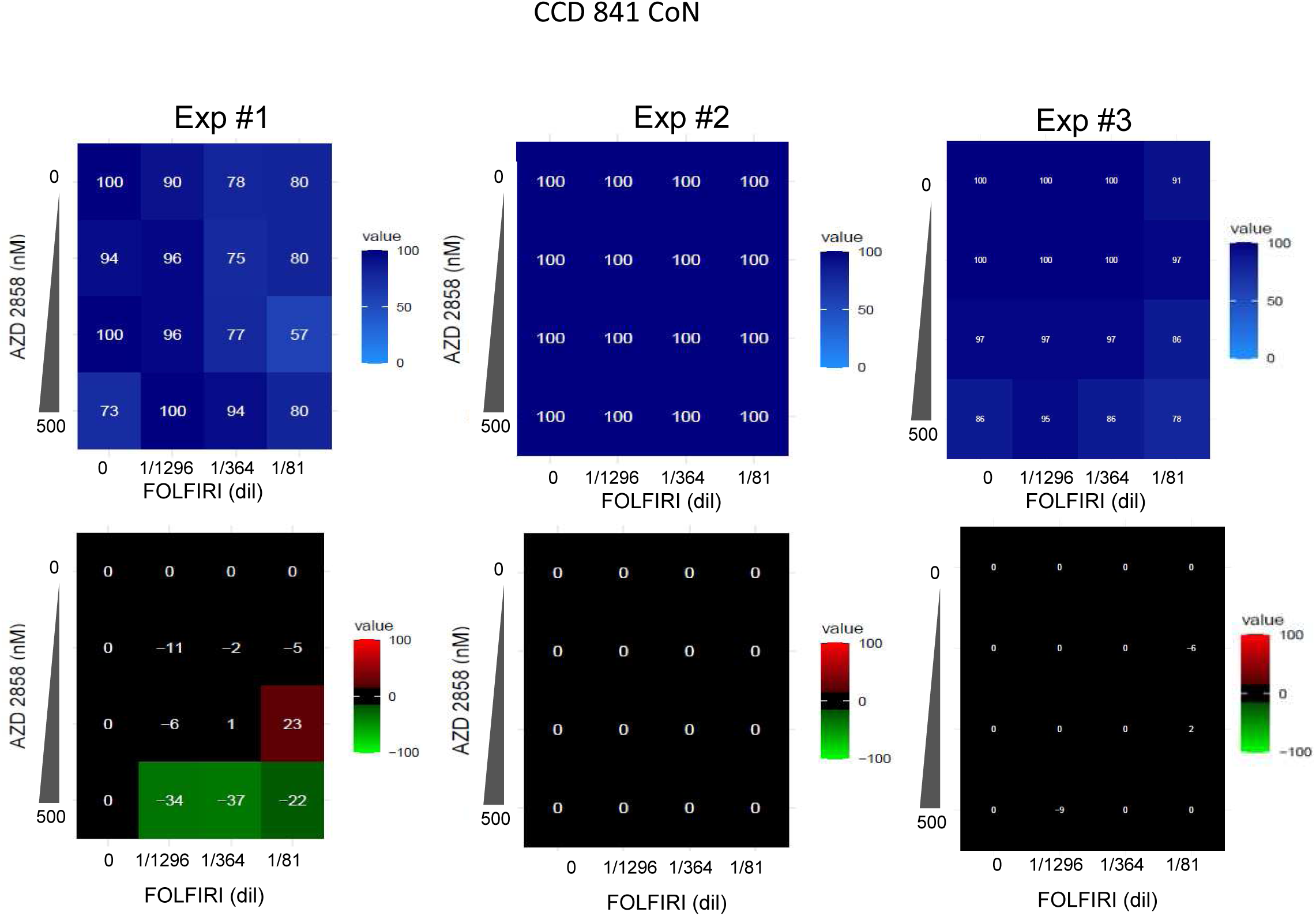
AZD2858 combined with FOLFIRI showed no synergistic effect on CCD 841 CoN cells. Untransformed colorectal cell lines, CCD 841 CoN were incubated with increasing concentrations of FOLFIRI (5−FU from 0.083 to 1.333 μM and SN−38 from 0.694 to 11.11 nM) and AZD2858 (from 7.8 to 500 nM). Cell viability was assessed with the SRB assay in 2D cultures to generate the **viability matrix** (blue). The **synergy matrices** (black, red and green) were calculated as described in Materials and Methods. Three independent experiments were shown.

**Table S1.** GSK-3 inhibitors in the TargetMol library. The Specificity column indicates the IC_50_ values available in the literature corresponding to 50% of the maximal concentration needed to inhibit the GSK-3 target (except for GSK3i XIII, where only the inhibition constant, Ki, value was available). The GSK-3β specificity column indicates the specificity toward this isoform, if available. If specificity was determined, but the GSK-3β isoform was not clearly specified, a question mark is used to indicate this uncertainty (for VP3.15 dihydrobromide and AT 7519 hydrochloride salt). ++++: < 1 nM; +++: between 1 nM and 10 nM; ++: between 10 nM and 40 nM; +: > 40 nM. Molecules that inhibited light-induced optoTopBP1 foci in the present screen are highlighted in red, and the z-score is indicated. The SN-38-induced Chk1 phosphorylation (pChk1) inhibition column indicates whether the potential GSK-3 inhibitors from the initial screen inhibit SN-38-induced Chk1 phosphorylation at S345 in HCT116 cells. N/A: Not Available.

| inhibitors of GSK3 in the bank | specificity | specificity to GSK-3β (IC50) | TopBP1 foci light induced inhibition | pChk1 SN-38 induced inhibition | source for IC50 data |
| --- | --- | --- | --- | --- | --- |
| 6-BIO | 5 nM ($\alpha/\beta$) | +++ | no | | selleckchem |
| 9-ING-41 | N/A | N/A | no |  | selleckchem |
| AR-A014418 | 38 nM ($\beta$) | ++ | no | | selleckchem |
| AT 7519 hydrochloride salt | 89 nM ($\alpha/\beta$?) | + (?) | yes (-3.4) | N/A | raybiotech |
| AT7519 | N/A | N/A | no |  | selleckchem |
| AZD1080 | 31 nM ($\beta$) 6.9 nM($\alpha$) | ++ | no | | selleckchem |
| AZD2858 | 5 nM ($\beta$) 0.9 nM ($\alpha$) | +++ | yes (-3.6) | yes | medchemexpress |
| Bikinin | N/A | N/A | no |  | selleckchem |
| BIO-acetoxime | 10 nM ($\alpha/\beta$) | ++ | yes (-3.7) | no | selleckchem |
| BRD0705 | 515 nM ($\beta$) | + | No | | selleckchem |
| CHIR98014 | 0.58 nM($\beta$) 0.65 nM ($\alpha$) | ++++ | yes (-2.8) | N/A | selleckchem |
| CHIR99021 | 6.7 nM ($\beta$) 10 nM ($\alpha$) | +++ | no | | selleckchem |
| CP21R7 | N/A | N/A | no |  | selleckchem |
| Ginsenoside Rg2 | N/A | N/A | no |  | N/A |
| GSK3i XIII | 24 nM (?) Ki | ++ | yes (-4.3) | N/A | sigma |
| GSK-3β inhibitor 1 | 4.19 nM | +++ | no | | selleckchem |
| IM12 | 53 nM ($\beta$) | + | no | | selleckchem |
| Indirubin | 600 nM ($\beta$) | + | no | | selleckchem |
| Indirubin-3'-monoxime | 190 nM | + | yes (-2.2) | N/A | tocris |
| KenPaullone | 230 nM ($\beta$) | + | no | | selleckchem |
| KY19382 | 10 nM($\beta$) | ++ | no | | medchemexpress+selleckchem |
| Lithium citrate tribasic tetrahydrate | N/A | N/A | no |  | N/A |
| LY2090314 | 0.9 nM ($\beta$) 1.5 nM ($\alpha$) | ++++ | yes (-3.4) | no | selleckchem |
| PHA767491 HCl | N/A | N/A | no |  | selleckchem |
| SB216763 | 34.3 nM ($\alpha/\beta$) | ++ | no | | selleckchem |
| SB415286 | 78 nM ($\alpha/\beta$) | + | no | | selleckchem |
| TBB | N/A | N/A | no |  | N/A |
| TDZD8 | 200 nM ($\beta$) | + | no | | selleckchem |
| Tideglusib | 60 nM ($\beta$) | + | no | | selleckchem |
| TWS119 | 30 nM ($\beta$) | ++ | yes (-2.8) | N/A | selleckchem |
| VP3.15 dihydrobromide | 880 nM ($\alpha/\beta$ ?) | + (?) | no | | medchemexpress |

**Table S2.**
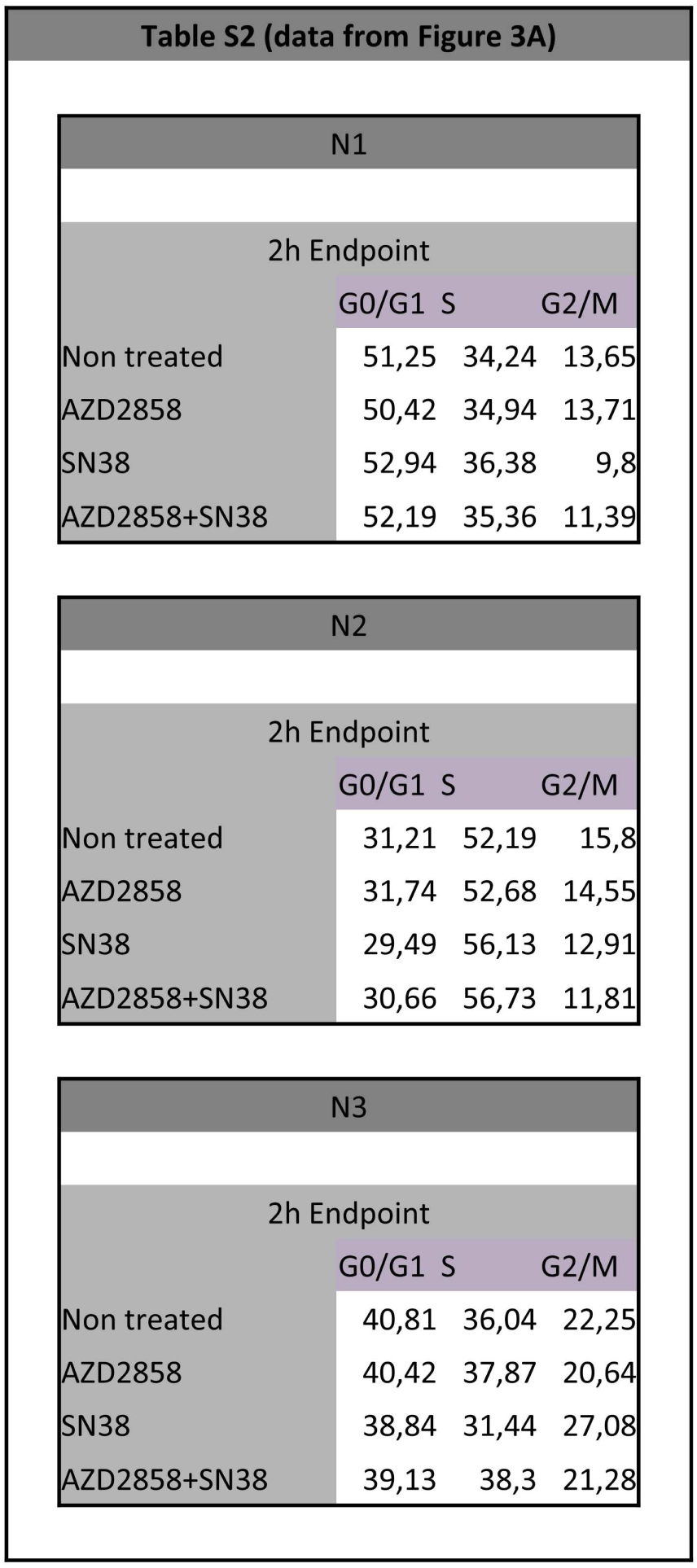
Three biological replicates results of flow cytometry experiments of the 2-hour condition (endpoint), including the representative one shown in Figure 3A. AZD2858 : 100 nM. SN-38 : 300 nM

**Table S3.** Three biological replicates results of flow cytometry experiments of the 6- and 12-hour condition (endpoint and pulse chase), including the representative one shown in Figure 3C and Figure S5C-D. AZD2858 : 100 nM. SN-38 : 300 nM

N1
|  | 6h Endpoint |  |  |  |
| --- | --- | --- | --- | --- |
|  | G0/G1 | S BrdU- | G2/M | S |
| Non treated | 26,73 | 0,66 | 12,54 | 59,61 |
| AZD2858 | 24 | 0,82 | 10,72 | 63,98 |
| SN38 | 41,7 | 12,19 | 17,93 | 27,12 |
| AZD2858+SN38 | 38,39 | 3,14 | 13,71 | 44,02 |

|  | 12h Endpoint |  |  |  |
| --- | --- | --- | --- | --- |
|  | G0/G1 | S BrdU- | G2/M | S |
| Non treated | 23,03 | 5,04 | 16,5 | 38,29 |
| AZD2858 | 24,39 | 1,04 | 17,6 | 51,38 |
| SN38 | 31,19 | 19,51 | 16,05 | 10,98 |
| AZD2858+SN38 | 35,96 | 4,06 | 13,65 | 44,08 |

|  | 6h Pulse Chase |  |  |  |  |  |
| --- | --- | --- | --- | --- | --- | --- |
|  | G0/G1 | SE | G2.b | G2 | S | N.G1 |
| Non treated | 41,41 | 23,12 | 2,66 | 17,46 | 9,38 | 5,24 |
| AZD2858 | 39,82 | 23,1 | 1,96 | 14,72 | 10,13 | 9,62 |
| SN38 | 44,99 | 0,63 | 19,13 | 8,2 | 24,56 | 1,49 |
| AZD2858+SN38 | 42,52 | 1,93 | 17,65 | 12,16 | 23,94 | 0,25 |

|  | 12h Pulse Chase |  |  |  |  |  |
| --- | --- | --- | --- | --- | --- | --- |
|  | G0/G1 | SE | G2.b | G2 | S | N.G1 |
| Non treated | 15,43 | 18,49 | 26,93 | 2,14 | 16,85 | 18,49 |
| AZD2858 | 21,39 | 21,94 | 19,8 | 1,51 | 22,01 | 11,3 |
| SN38 | 43,22 | 1,35 | 18,17 | 11,43 | 20,32 | 3,1 |
| AZD2858+SN38 | 47,17 | 7,55 | 12,59 | 15,28 | 15,11 | 1,11 |

N2
|  | 6h Endpoint |  |  |  |
| --- | --- | --- | --- | --- |
|  | G0/G1 | S BrdU- | G2/M | S |
| Non treated | 22,21 | 0,7 | 12,5 | 64,17 |
| AZD2858 | 11,53 | 0,82 | 12,19 | 75,17 |
| SN38 | 59,08 | 14,88 | 10,4 | 15,25 |
| AZD2858+SN38 | 45,1 | 2,66 | 10,7 | 41,01 |

|  | 12h Endpoint |  |  |  |
| --- | --- | --- | --- | --- |
|  | G0/G1 | S BrdU- | G2/M | S |
| Non treated | 30,87 | 0,8 | 19,05 | 48,92 |
| AZD2858 | 31,65 | 0,99 | 18,29 | 48,74 |
| SN38 | 59,5 | 9,55 | 9,95 | 20,24 |
| AZD2858+SN38 | 53,73 | 2,93 | 9,88 | 32,79 |

|  | 6h Pulse Chase |  |  |  |  |  |
| --- | --- | --- | --- | --- | --- | --- |
|  | G0/G1 | SE | G2.b | G2 | S | N.G1 |
| Non treated | 18,77 | 35,33 | 1,41 | 25,03 | 10,7 | 8,41 |
| AZD2858 | 16,63 | 38,28 | 1,01 | 25,07 | 10,05 | 8,73 |
| SN38 | 46,73 | 0,51 | 7,57 | 4,25 | 39,64 | 0,87 |
| AZD2858+SN38 | 49,32 | 2,52 | 6,23 | 7,01 | 33,43 | 0,86 |

|  | 12h Pulse Chase |  |  |  |  |  |
| --- | --- | --- | --- | --- | --- | --- |
|  | G0/G1 | SE | G2.b | G2 | S | N.G1 |
| Non treated | 12,13 | 26,94 | 22,42 | 2,41 | 14,36 | 21,09 |
| AZD2858 | 13,02 | 15,95 | 28,57 | 2,48 | 23,19 | 16,34 |
| SN38 | 58,87 | 2,61 | 5,42 | 4,2 | 26,14 | 1,59 |
| AZD2858+SN38 | 47,98 | 13,04 | 5,55 | 10,97 | 21,11 | 0,5 |

N3
|  | 6h Endpoint |  |  |  |
| --- | --- | --- | --- | --- |
|  | G0/G1 | S BrdU- | G2/M | S |
| Non treated | 24,21 | 0,59 | 14,64 | 60,03 |
| AZD2858 | 22,85 | 0,6 | 14,02 | 61,97 |
| SN38 | 54,22 | 18,78 | 13,08 | 13,36 |
| AZD2858+SN38 | 40,45 | 2,53 | 10,43 | 45,72 |

|  | 12h Endpoint |  |  |  |
| --- | --- | --- | --- | --- |
|  | G0/G1 | S BrdU- | G2/M | S |
| Non treated | 26,13 | 0,61 | 17,01 | 54,7 |
| AZD2858 | 27,47 | 0,54 | 13,65 | 57,1 |
| SN38 | 54,9 | 15,27 | 12,37 | 12,11 |
| AZD2858+SN38 | 48,86 | 3,3 | 7,17 | 38,84 |

|  | 6h Pulse Chase |  |  |  |  |  |
| --- | --- | --- | --- | --- | --- | --- |
|  | G0/G1 | SE | G2.b | G2 | S | N.G1 |
| Non treated | 27,67 | 31,74 | 1,26 | 14,37 | 17,75 | 6,84 |
| AZD2858 | 24,79 | 33,13 | 0,88 | 16,57 | 15,52 | 8,69 |
| SN38 | 45,67 | 2,38 | 8,6 | 4,79 | 37,39 | 0,87 |
| AZD2858+SN38 | 45,77 | 4,58 | 7,66 | 7,11 | 34,1 | 0,25 |

|  | 12h Pulse Chase |  |  |  |  |  |
| --- | --- | --- | --- | --- | --- | --- |
|  | G0/G1 | SE | G2.b | G2 | S | N.G1 |
| Non treated | 14,04 | 13,25 | 16,65 | 1,99 | 22,64 | 30,91 |
| AZD2858 | 19,36 | 11,6 | 16,19 | 2,13 | 27,62 | 22,3 |
| SN38 | 49,88 | 1,81 | 7,52 | 5,41 | 31,15 | 2,94 |
| AZD2858+SN38 | 51,79 | 5,64 | 5,64 | 8,74 | 25,98 | 1,03 |

**Table S4.** Results of the linear regression analysis(Figure 4A-B). NT: Non−Treated. A: AZD2858. S: SN−38. F: FOLFIRI (5−FU + SN−38). The p−values of interest from Figure 4B are indicated in red.

Estimation of effect (amplitude and p-value) by linear modeling: Effect of experimental condition (compared to "NT") :
| Effect on percentage of |  |  |
| --- | --- | --- |
| Condition | subG1 | p-value |
| A | -0.05329 | 0.99220 |
| AF | +56.43905 | 9.19e-07 |
| AS | +46.09911 | 5.79e-06 |
| F | +17.95105 | 0.00706 |
| S | +9.60998 | 0.10089 |

Effect of experimental condition (compared to "S") :
| Effect on percentage of |  |  |
| --- | --- | --- |
| Condition | subG1 | p-value |
| NT | -9.610 | 0.10089 |
| A | -9.663 | 0.09926 |
| AF | +46.829 | 5.03e-06 |
| AS | +36.489 | 4.40e-05 |
| F | +8.341 | 0.14786 |

Effect of experimental condition (compared to "F") :
| Effect on percentage of |  |  |
| --- | --- | --- |
| Condition | subG1 | p-value |
| S | -8.341 | 0.147855 |
| NT | -17.951 | 0.007057 |
| A | -18.004 | 0.006939 |
| AF | +38.488 | 2.8e-05 |
| AS | +28.148 | 0.000351 |

Effect of replicate identity (compared to replicate #1) :
| Effect on percentage of |  |  |
| --- | --- | --- |
| Replicate | subG1 | p-value |
| 2 | +12.45762 | 0.00784 |
| 3 | +11.50860 | 0.01204 |

**Table S5.** FOLFIRI concentration ranges used for the synergy matrix assays. All the dilutions are based on the dilution 1/1 corresponding to 12 µM 5-fluorouracil (5-FU) and 100 nM SN-38.

| FOLFIRI concentration range<br>(dilution 1/1 = 12 µM 5-FU + 100 nM SN-38) |  |  |  |  |  |  |
| --- | --- | --- | --- | --- | --- | --- |
| HCT116 (2D & 3D) | **FOLFIRI (dilution factor)** | **1296** | **648** | **324** | **162** | **81** |
|  | 5-FU (µM) | 0,009 | 0,019 | 0,037 | 0,074 | 0,148 |
|  | SN-38 (nM) | 0,077 | 0,154 | 0,309 | 0,617 | 1,235 |
| CT26 (2D & 3D) | **FOLFIRI (dilution factor)** | **144** | **72** | **36** | **18** | **9** |
|  | 5-FU (µM) | 0,083 | 0,167 | 0,333 | 0,667 | 1,333 |
|  | SN-38 (nM) | 0,694 | 1,389 | 2,778 | 5,556 | 11,111 |
| HT29 (2D) | **FOLFIRI (dilution factor)** | **432** | **216** | **108** | **54** | **27** |
|  | 5-FU (µM) | 0,028 | 0,056 | 0,111 | 0,222 | 0,444 |
|  | SN-38 (nM) | 0,231 | 0,463 | 0,926 | 1,852 | 3,704 |
| SW620 (2D) | **FOLFIRI (dilution factor)** | **864** | **432** | **216** | **108** | **54** |
|  | 5-FU (µM) | 0,014 | 0,028 | 0,056 | 0,111 | 0,222 |
|  | SN-38 (nM) | 0,116 | 0,231 | 0,463 | 0,926 | 1,852 |
| SW480 (2D) | **FOLFIRI (dilution factor)** | **864** | **432** | **216** | **108** | **54** |
|  | 5-FU (µM) | 0,014 | 0,028 | 0,056 | 0,111 | 0,222 |
|  | SN-38 (nM) | 0,116 | 0,231 | 0,463 | 0,926 | 1,852 |
| HCT116-SN6 (2D)<br>HCT116-SN50 (2D) | **FOLFIRI (dilution factor)** | **32** | **16** | **8** | **4** | **2** |
|  | 5-FU (µM) | 0,375 | 0,750 | 1,500 | 3,000 | 6,000 |
|  | SN-38 (nM) | 3,125 | 6,250 | 12,500 | 25,000 | 50,000 |
| HCT116-SN6 (3D) | **FOLFIRI (dilution factor)** | **192** | **96** | **48** | **24** | **12** |
|  | 5-FU (µM) | 0,063 | 0,125 | 0,250 | 0,500 | 1,000 |
|  | SN-38 (nM) | 0,521 | 1,042 | 2,083 | 4,167 | 8,333 |
| HCT116-SN50 (3D) | **FOLFIRI (dilution factor)** | **48** | **24** | **12** | **6** | **3** |
|  | 5-FU (µM) | 0,250 | 0,500 | 1,000 | 2,000 | 4,000 |
|  | SN-38 (nM) | 2,083 | 4,167 | 8,333 | 16,667 | 33,333 |

**Table S6.** References of the antibodies used in the study. WB, western blotting, IF, immunofluorescence analyses.

| Target | Compagny | Reference | Species | Dilution Celigo | Dilution WB | Dilution IF |
| --- | --- | --- | --- | --- | --- | --- |
| H3 | Cell Signaling Technology | #4499 | Rabbit |  | 1/1000 |  |
| α-Tubulin | Sigma | #T5168 | Mouse | | 1/10000 | |
| 53BP1 | Abcam | #ab175933 | Rabbit |  |  | 1/100 |
| ATR | Bethyl | #A300-137A | Rabbit |  | 1/1000 |  |
| Chk1 | Santa Cruz Biotechnology | #sc-8408 | Mouse |  | 1/1000 |  |
| Chk2 | Milipore | #05-649 | Mouse |  | 1/1000 |  |
| Cleaved caspase-3 | Cell Signaling Technology | #9661 | Rabbit |  | 1/1000 |  |
| GSK-3β | Cell Signaling Technology | #9832 | Mouse | | 1/1000 | |
| GSK-3β (S9) | Cell Signaling Technology | #9336 | Rabbit | | 1/1000 | |
| H2B | Abcam | #ab1790 | Rabbit |  | 1/2500 |  |
| pATM (T1981) | Santa Cruz Biotechnology | #sc-47739 | Mouse | 1/100 |  |  |
| pATM (T1981) | Rockland | #200-301-40 | Mouse |  | 1/1000 |  |
| pChk1 (S345) | Cell Signaling Technology | #2348 | Rabbit | 1/50 | 1/1000 |  |
| pChk2 (T68) | Cell signaling Technology | #2661 | Rabbit | 1/50 | 1/1000 |  |
| PARP1 | Santa Cruz Biotechnology | #sc-8007 | Mouse |  | 1/1000 |  |
| PML | Santa Cruz Biotechnology | #sc-966 | Mouse |  |  | 1/100 |
| Pol $\epsilon$ | Kind gift from Shou Waga | #N/A | Rabbit | | 1/1000 | |
| pRPA32 (S33) | Abcam | #ab2118877 | Rabbit |  | 1/1000 |  |
| RPA32 | Abcam | #ab2175 | Mouse |  | 1/1000 |  |
| RPA34 | Kind gift from Marcel Méchali |  | Rabbit |  | 1/1000 |  |
| TopBP1 | Santa Cruz | #sc-271043 | Mouse |  |  | 1/100 |
| TopBP1 | Euromedex | #A300-111A | Rabbit |  | 1/1000 |  |
| TopBP1 | Kind gift from Larry Karnitz |  | Rabbit |  | 1/1000 |  |
| Vinculin | Merck | #V9131 | Mouse |  | 1/10000 |  |
| γH2AX (S139) | Cell Signaling Technology | #9718S | Rabbit | | 1/1000 | |
| γH2AX (S139) | Abcam | #ab1195189 | Rabbit | 1/1000 | | |
| Goat anti-rabbit IgG (H+L) AF568 | Thermo Fisher Scientific | #A11011 | Secondary |  | 1/5000 |  |
| Goat anti-mouse IgG (H+L) AF488 | Thermo Fisher Scientific | #A11029 | Secondary |  | 1/5000 |  |

**Manuscript number:** RC-2024-02687

**Corresponding author(s):** Jihane Basbous

## 1. General Statements [optional]

*This section is optional. Insert here any general statements you wish to make about the goal of the study or about the reviews*.

## 2. Point-by-point description of the revisions

Reviewer #1 (Evidence, reproducibility and clarity (Required)):

### Summary

Laura Morano and colleagues have performed a screen to identify compounds that interfere with the formation of TopBP1 condensates. TopBP1 plays a crucial role in the DNA damage response, and specifically the activation of ATR. They found that the GSK-3b inhibitor AZD2858 reduced the formation of TopBP1 condensates and activation of ATR and its downstream target CHK1 in colorectal cancer cell lines treated with the clinically relevant irinotecan active metabolite SN-38. This inhibition of TopBP1 condensates by AZD2858 was independent from its effect on GSK-3b enzymatic activity. Mechanistically, they show that AZD2858 thus can interfere with intra-S-phase checkpoint signaling, resulting in enhanced cytostatic and cytotoxic effects of SN-38 (or SN-38+Fluoracil aka FOLFIRI) in vitro in colorectal carcinoma cell lines.

### Major comments

Overall the work is rigorous and the main conclusions are convincing. However, they only show the effects of their combination treatments on colorectal cancer cell lines. I’m worried that blocking the formation of TopB1 condensates will also be detrimental in non-transformed cells. Furthermore it is somewhat disappointing that it remains unclear how AZD2858 blocks self-assembly of TopBP1 condensates, although I understand that unraveling this would be complex and somewhat out-of-reach for now.

We appreciate your feedback and fully recognize the importance of understanding how AZD2858 blocks the assembly of TopBP1 condensates. While we understand your disappointment, addressing this question remains a key focus for us. Keeping in mind that unravelling such a mechanism in vitro or in vivo is rather challenging, we have consulted an expert who has made efforts to predict the potential docking sites of AZD2858 on TopBP1, which may provide valuable insights for future experimental investigations. Using an AlphaFold model (no crystal or cryo-EM structure available) and looking for suitable pockets or cavities in which AZD2858 could bind, the analyses, though requiring cautious interpretation, suggested that AZD2858 may target the BRCT1 and BRCT8 domains (as shown below, two pockets n°1 and 7 with sufficient volume and surrounded by b-sheets structures like other GSK3 inhibitor) of TopBP1. However, these are preliminary results that require further exploration and experimental validation to confirm their significance and mechanistic implications.

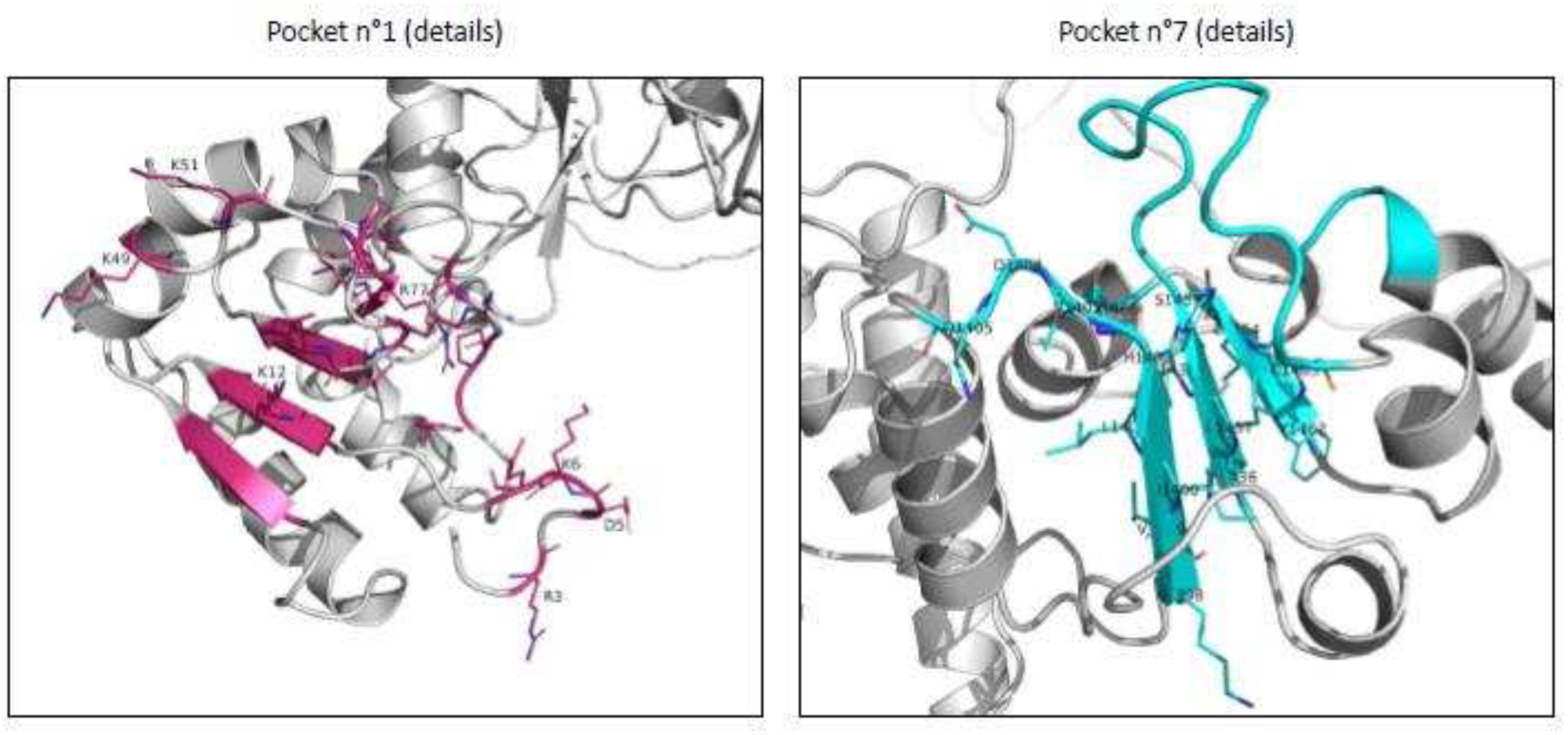

Here are some specific points for improvement:

1) The authors conclude that “These data supports [sic] the feasibility of targeting condensates formed in response to DNA damage to improve chemotherapy-based cancer treatments”. To support this conclusion the authors need to show that proliferating non-transformed cells (e.g. primary cell cultures or organoids) can tolerate the combination of AZD2858 + SN-38 (or FOLFIRI) better than colorectal cancer cells.

We would like to thank the reviewer for this vital suggestion to prove that this combination is effective on tumor cells and not very toxic on healthy cells. We therefore used a healthy colon cell line (CCD841) and tested the efficacy of each treatment alone (FOLFIRI and AZD2858) as well as the combination FOLFIRI+AZD2858. We compared the results obtained in the CCD841 cell line with those obtained in the HCT116 colorectal cancer cell line. The results presented below show not only that each treatment alone is much less effective on CCD841 lines, but also that the combination is not synergistic.

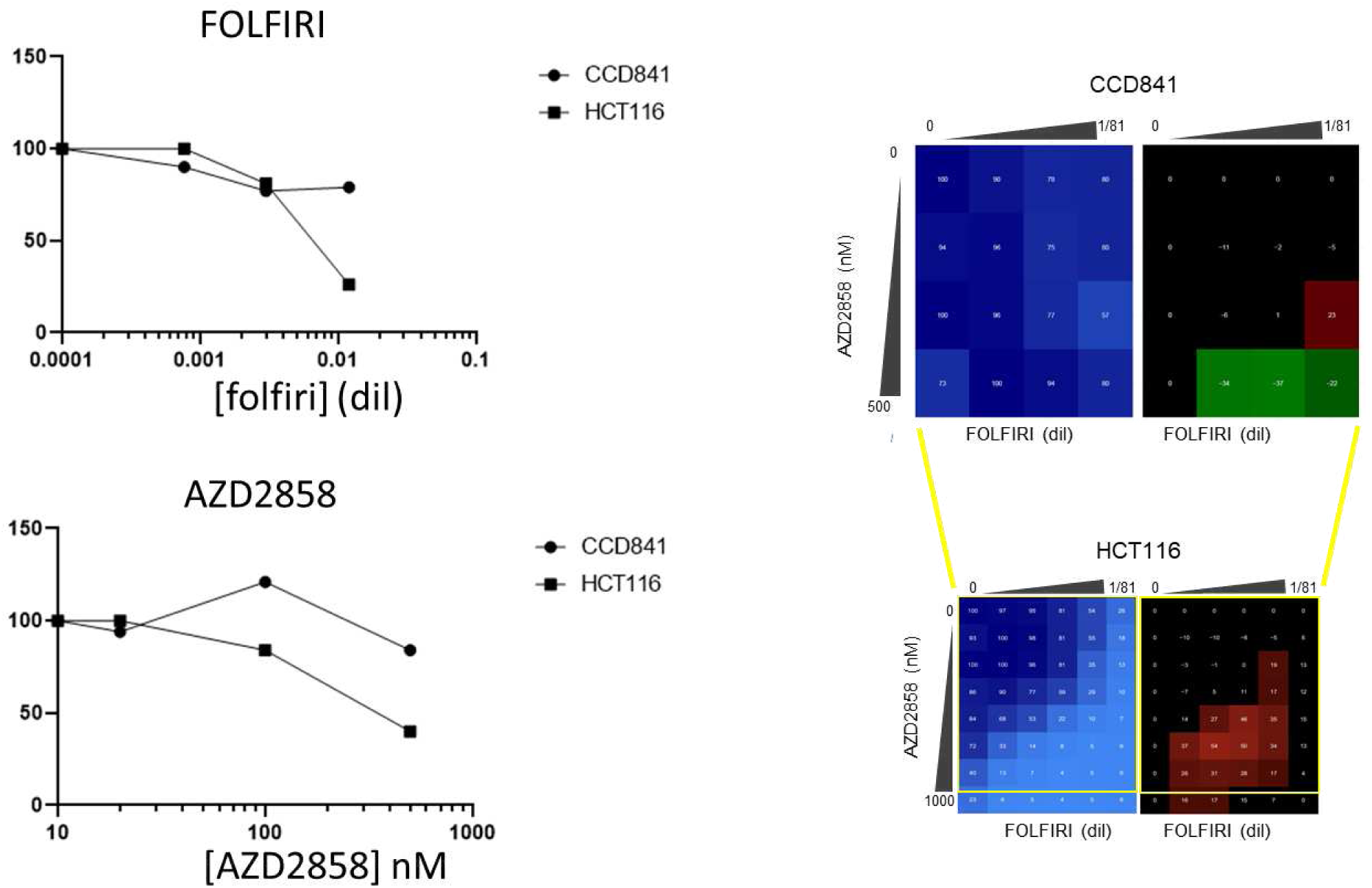

2) Page 19 “This suggests that the combination… arrests the cell cycle before mitosis in a DNA-PKsc-dependent manner.” I find the remark that this arrest would be DNA-PKcs-dependent too speculative. I suppose that the authors base this claim on reference 55 but if they want to support this claim they need to prove this by adding DNA-PKcs inhibitors to their treated cells.

Thank you for your thoughtful comment. We agree with the reviewer that claiming the G2/M arrest is DNA-PKcs-dependent without direct experimental evidence is speculative. While we initially based this hypothesis on reference 55, we acknowledge that further experiments, such as the use of DNA-PKcs inhibitors, would be necessary to robustly support this claim.

Given that this observation was intended as a potential explanation for the G2/M arrest observed at 6 and 12 hours of treatment with AZD2858 + SN-38 (compared to SN-38 alone), and considering that exploring this pathway is not the primary focus of our study, we have decided to remove this hypothesis from both the figure and the text to avoid any ambiguity.

We appreciate the reviewer’s input and will consider investigating this pathway in future studies.

3) When discussing Figure S5B the authors claim that SN-38 + AZD2858 progressively increases the fractions of BrdU positive cells, but this is not supported by statistical analysis.

The fractions are still very small, so I would like to see statistics on these data. Alternatively, the authors could take out this conclusion.

Thank you for your valuable comment. In response, we have conducted a statistical analysis (Mann-Whitney test) on the data, and the results have been added to **Figure S5C** for the 6-hour time point and **Figure S5D** for the 12-hour time point, based on three independent biological replicates. We hope this provides the necessary clarification.

### Minor comments

- Page 5 Materials and methods - Cell culture. Last sentence “Add in what medium you cultured them” looks like an internal review remark and should probably be removed?

We apologize for this oversight. The medium has now been specified, and the sentence has been removed.

- The numbers in all the synergy matrices (in white font) are extremely small and virtually unreadable, and visually distracting. I recommend taking these out altogether.

We believe that the reduction in figure quality may be due to the PDF compression, which affected the resolution of the figures. We are happy to provide high-resolution versions of the figures separately for clarity. If the issue persists even with the higher resolution, we will consider removing the numbers, as suggested.

- The legends of the synergy matrices (for example Fig 1D, 4E, 5, 6) are often extremely small, making it difficult to understand them intuitively. Please enlarge them and label them more clearly, and use larger fonts. In the legend of Figure 5D,E a green matrix indicating % live cells is mentioned but I don’t see it. Do they mean the grey matrix?

We have enlarged the figure legends and will provide high-resolution versions of the figures to ensure all details are clearly readable. Regarding **Figure 5D,E**: we acknowledge that the color may appear differently (more green or gray) depending on the display or printer settings. To avoid any confusion, we have corrected the legend to specify that the color in question is khaki, rather than green. Moreover, following suggestions of the reviewer #2, these figures have been respectively moved to **Figure S6B and S6C.**

- Figure S2. Perhaps I misunderstand the PML body experiment but the authors seem to use PML body formation to support their idea that AZD2858 blocks TopBP1 condensate formation and not just any condensate formation. However, if this is the case they would need a proper positive control, i.e. an additional experimental condition in which they do see PLM bodies.

Arsenic is a well-known positive control for experiments involving PML bodies due to its ability to induce specific responses in PML proteins and modify PML nuclear bodies (NBs) structure and function (Jaffray et al., 2023, JCB ; Zhu et al., 1997, PNAS). Thus, we used Arsenic as a positive control and observed a significant increase in PML NBs vs the other conditions (Kruskal-Wallis test) as indicated below. We thus implemented the results in the corresponding figure S2B and text.

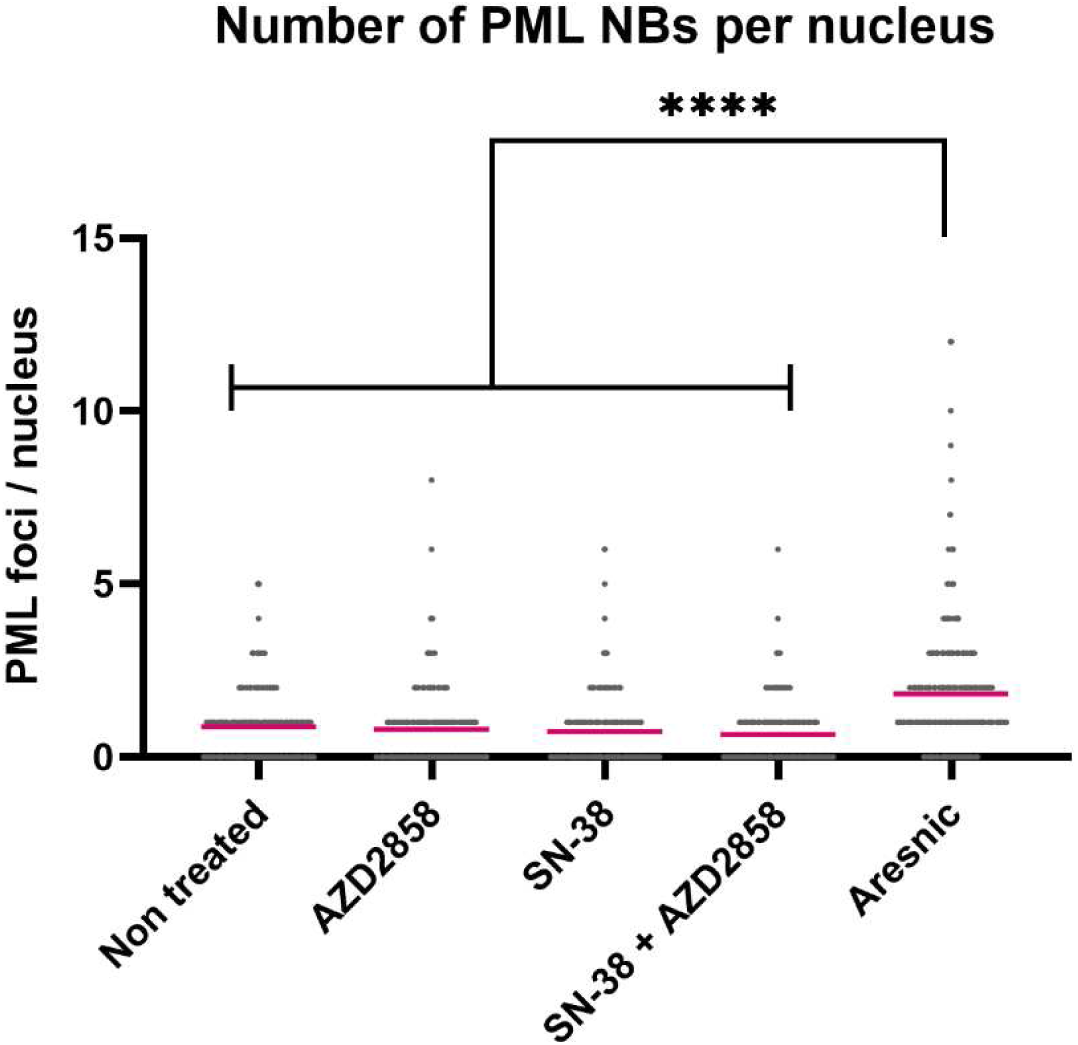

**Legend :**PML condensates were tested after 2 h of incubation. AZD2858 : 100nM ; SN-38 : 300nM ; Arsenic : 6µM. ****: p<0.0001 (Kruskal-Wallis test).

- The quantification of the flow cytometry data needs to be clarified. I find it strange that in the figures (for example Figure 3A and 3C) representative examples are shown of apparently 3 replicates, and that the percentages shown in these examples are then the given in the text as the overall numbers; for example on page 18 “…BrdU incorporation increased from 16.11% (SN-38 alone) to 41.83% (combination)…”. This type of description is done in multiple places in the Results section and is confusing. It would be clearer if the authors show proper quantifications (mean +/-sem) of the percentages of (the relevant) gated populations. Besides, I don’t think it make a lot of sense to mention in the text the percentages with 2 decimals behind the comma.

This suggests a level of precision that does not seem justified in flow cytometry data. Finally, all flow cytometry plots look visually very busy and all the text is crammed in with really small fonts. Cleaning them up and enlarging the fonts of the remaining text/numbers would really improve the readability of the figures.

Thank you for your helpful comments. We understand your concern regarding the flow cytometry quantification. Indeed, the percentages presented in the figures are derived from representative replicates, and we acknowledge that this presentation could be confusing. To address this, we have included a table summarizing the data from all replicates to improve readability [**Table S2 and S3** in the new version]. Second, we specified in the text that the data are representative biological replicates when needed. Third, we have performed statistical analyses on the three replicates when necessary, as shown in Supplementary **Figure S5C-F** in the new version. The text has been revised to reflect the correct statistical interpretation.

Regarding the use of two decimal, we are unable to remove them due to limitations in the software (Kaluza) used for flow cytometry analysis. However, we agree that this level of precision may not be warranted, and we have revised the text where appropriate to reduce confusion.

- In Figure 5G the authors show that FOLFIRI + AZD2858 are synergistic in two SN-38-resistant cell lines. They conclude that this combination may overcome drug resistance. But tried to figure out the used FOLFIRI concentrations used in these cell lines and they still seem far higher than the SN-38-sensitive HCT116 cell lines, so I would like to see a bit more nuance in their interpretation. I think overcoming drug resistance is an overstatement, and perhaps alleviating would be a better term

Thank you for highlighting this important point; we have adjusted the text accordingly.

-The legend in Table S2 refers to Figure 5A-B; this should be Figure 4A-B. Thank you, this has been corrected and Table S2 is now moved to Table S4.

Reviewer #1 (Significance (Required)):

SECTION B - Significance

========================

The finding that AZD2858 block TOPbp1 condensate formation via a pleiotropic effect of this compound is interesting and convincing. To my best knowledge it’s a novel finding which is interesting to the potential target audience mentioned below. Their findings that inhibition of TOPbp1 condensation and ATR signaling via AZD2858 may synergize with FOLFIRI therapy in colorectal cancer cells are still very preliminary, because the effects on non-cancerous cells are not tested.

Researchers involved in early cancer drug discovery and cell biologists studying DNA damage responses in cancer cells seem to me typical audience interested and influenced by this paper.

I’m a cell biologist studying cell cycle fate decisions, and adaptation of cancer cells & stem cells to (drug-induced) stress. My expertise aligns well with the work presented throughout this paper.

Reviewer #2 (Evidence, reproducibility and clarity (Required)):

The authors have extended their previous research to develop TOPBP1 as a potential drug target for colorectal cancer by inhibiting its condensation. Utilizing an optogenetic approach, they identified the small molecule AZD2858, which inhibits TOPBP1 condensation and works synergistically with first-line chemotherapy to suppress colorectal cancer cell growth. The authors investigated the mechanism and discovered that disrupting TOPBP1 assembly inhibits the ATR/Chk1 signaling pathway, leading to increased DNA damage and apoptosis, even in drug-resistant colorectal cancer cell lines. Addressing the following concerns would enhance clarity and further in vivo work may improve significance:

1. How does the optogenetic method for inducing condensates compare to the DNA damage induction mechanism?

Optogenetics provides a versatile and precise approach for controlling the condensation of scaffold proteins in both space and time. This method enables us to study the role of biomolecular condensates with minute-scale resolution, separating their formation from potentially confounding upstream events, such as DNA damage, and providing valuable insights into their specific function. Importantly, based on our previous publications on TopBP1 or SLX4 optogenetic condensates, we have substantial evidence indicating that light-induced condensates closely mimic those formed in response to DNA damage:

### Functional similarity

Optogenetic condensates recapitulate endogenous condensates formed upon exposure of the cells of DNA damaging agents, and include most known partner proteins involved in the DNA damage response. It was shown for light induced-TopBP1 and SLX4 condensates (1–3).

### Dynamic reversibility

Optogenetic condensates and DNA damage induced condensates are both dynamic and reversible. They dissolve within 15 minutes of light deactivation or after removal of the damaging agent (1,3).

### Chromatin association

Both optogenetic and DNA damage-induced condensates are bound to chromatin or localized at sites of DNA damage (3).

### Regulation

Both types of condensates are regulated similarly, with their formation triggered by the same signaling pathways. ATR basal activity drives the nucleation of opto-TopBP1 condensates and endogenous TopBP1 structures upon light exposure (1). Likewise, sumoylation modifications regulate the formation of opto-SLX4 condensates and endogenous SLX4 condensates (3).

### Structurally

Using super-resolution imaging by stimulation-emission-depletion (STED) microscopy, we observed that endogenous SLX4 nanocondensates formed globular clusters that were indistinguishable from recombinant light induced SLX4 condensates (1,3).

## 2. Why wasn’t the initial screen conducted on the HCT116-SN50 resistant cell line?

Thank you for raising this important question, which we also considered at the outset of the project. After careful consideration, we decided to use the HCT116 WT cells in order to obtain initial data from an unmodified cell line. It is worth mentioning that HCT116-SN50 cells exhibit slower proliferation compared to WT cells, and they also express an efflux pump capable of pumping out SN38. We were concerned that these factors might interfere with the optogenetic assay, which is why we chose to perform the screen using the WT HCT116 cells.

## 3. The labels in Fig. 1D are difficult to recognize

This issue was also raised by Reviewer #1. We suspect that the PDF conversion may have reduced the resolution of the figures, so we will provide them separately in high resolution. In addition, we have increased the size of some labels to improve their clarity.

## 4. The selected cell image in Fig. 2A for SN-38 seems over-representative; unselected cells appear similar to other groups. Why does AZD2858 itself induce TopBP1 condensates in the plot, yet this is not evident in the images?

Thank you for your comment; we have updated the figure with a more representative image. We indeed observe that AZD2858 alone induces a slight increase in TopBP1 condensates. However, this increase did not lead to the activation of the ATR/Chk1 signaling pathway, as shown by the Western blot data presented in Fig. 2B. In addition, AZD2858 specifically prevents the formation of TopBP1 condensates induced by SN38 treatment, and the level of TopBP1 condensates does not return to the basal levels observed in untreated cells, but rather to those observed with AZD2858 treatment. During the 2-hour AZD2858 treatment, the progression of replication forks was unaffected (Fig. 3A and 3B). However, when AZD2858 was added alone to the Xenopus egg extracts, there was increased recruitment of TopBP1 to the chromatin (Fig. 2E). This result suggests that AZD2858 alone can induce the assembly of TopBP1 on chromatin to initiate DNA replication (a well-established role of TopBP1), but the number and concentration of TopBP1 molecules did not reach levels sufficient to activate the ATR/Chk1 pathway.

## 5. In Fig. 3A, despite the drastic change in the FACS plot shape, the quantifications appear quite similar

Thank you for this insightful observation. The gates for the S phase were intentionally set wider to avoid biasing the results and inadvertently excluding the population that incorporates BrdU weakly (but still incorporates it) in the SN-38 only condition. As a result, the percentage of cells within this gate remains similar, even though the overall shape of the FACS plot changes, reflecting a shift in the distribution of BrdU incorporation. This point has now been clarified in the legend of the **Figure 3A**.

This effect can also be attributed to the relatively short treatment time (2 hours), which captures early changes in DNA synthesis. The effect becomes more pronounced at later time points, as shown in **Figure 3C**. For example, after 6 hours of treatment, the percentage of BrdU-positive cells increases from 15% with SN-38 alone to 41% with the AZD2858 combination, demonstrating a clearer impact on DNA synthesis. A graph summarizing the statistical analysis has been added to **Figure S5C** for the 6-hour time point and **Figure S5D** for the 12-hour time point, based on data from three independent biological replicates.

## 6. The results section is imbalanced; Figs. 5 and 6 could be combined into one figure

We have combined Figures 5 and 6 into a single figure to optimize the presentation of results. To avoid overloading the new figure, some of the data have been moved to supplementary figures, ensuring the main figure remains clear and focused.

## 7. An in vivo study is anticipated to assess the drug’s efficacy

Although AZD2858 was developed a few years ago, there is a limited amount of in vivo data available, which led us to consider potential issues related to the drug’s biodistribution or its pharmacokinetics (PK). Despite these concerns, we proceeded with preliminary in vivo studies, testing various diluents and injection routes for AZD2858. However, we observed that the compound was not effective in vivo. Given the strong synergistic effects observed in vitro, we concluded that AZD2858 was likely not being distributed properly in the mice. As a result, we have decided to conduct a more detailed investigation into the pharmacokinetics (PK), pharmacodynamics (PD), and absorption, distribution, metabolism, and excretion (ADME) of AZD2858 to better understand its in vivo behavior and efficacy. Therefore, the in vivo evaluation of AZD2858 will be addressed in a separate study specifically focused on this aspect.

Reviewer #2 (Significance (Required)):

Addressing the stated concerns would enhance clarity and further in vivo work may improve significance.

Reviewer #3 (Evidence, reproducibility and clarity (Required)):

### Summary

In 2021 (PMID: 33503405) and 2024 (PMID: 38578830) Constantinou and colleagues published two elegant papers in which they demonstrated that the Topbp1 checkpoint adaptor protein could assemble into mesoscale phase-separated condensates that were essential to amplify activation of the PIKK, ATR, and its downstream effector kinase, Chk1, during DNA damage signalling. A key tool that made these studies possible was the use of a chimeric Topbp1 protein bearing a cryptochrome domain, Cry2, which triggered condensation of the chimeric Topbp1 protein, and thus activation of ATR and Chk1, in response to irradiation with blue light without the myriad complications associated with actually exposing cells to DNA damage.

In this current report Morano and co-workers utilise the same optogenetic Topbp1 system to investigate a different question, namely whether Topbp1 phase-condensation can be inhibited pharmacologically to manipulate downstream ATR-Chk1 signalling. This is of interest, as the therapeutic potential of the ATR-Chk1 pathway is an area of active investigation, albeit generally using more conventional kinase inhibitor approaches.

The starting point is a high throughput screen of 4730 existing or candidate small molecule anti-cancer drugs for compounds capable of inhibiting the condensation of the Topbp1-Cry2-mCherry reporter molecule in vivo. A surprisingly large number of putative hits (>300) were recorded, from which 131 of the most potent were selected for secondary screening using activation of Chk1 in response to DNA damage induced by SN-38, a topoisomerase inhibitor, as a surrogate marker for Topbp1 condensation. From this the 10 most potent compounds were tested for interactions with a clinically used combination of SN-38 and 5-FU (FOLFIRI) in terms of cytotoxicity in HCT116 cells. The compound that synergised most potently with FOLFIRI, the GSK3-beta inhibitor drug AZD2858, was selected for all subsequent experiments.

AZD2858 is shown to suppress the formation of Topbp1 (endogenous) condensates in cells exposed to SN-38, and to inhibit activation of Chk1 without interfering with activation of ATM or other endpoints of damage signalling such as formation of gamma-H2AX or activation of Chk2 (generally considered to be downstream of ATM). AZD2858 therefore seems to selectively inhibit the Topbp1-ATR-Chk1 pathway without interfering with parallel branches of the DNA damage signalling system, consistent with Topbp1 condensation being the primary target.

Importantly, neither siRNA depletion of GSK3-beta, or other GSK3-beta inhibitors were able to recapitulate this effect, suggesting it was a specific non-canonical effect of AZD2858 and not a consequence of GSK3-beta inhibition per se.

To understand the basis for synergism between AZD2858 and SN-38 in terms of cell killing, the effect of AZD2858 on the replication checkpoint was assessed. This is a response, mediated via ATR-Chk1, that modulates replication origin firing and fork progression in S-phase cell under conditions of DNA damage or when replication is impeded. SN-38 treatment of HCT116 cells markedly suppresses DNA replication, however this was partially reversed by co-treatment with AZD2858, consistent with the failure to activate ATR-Chk1 conferring a defect in replication checkpoint function.

Figures 4 and 5 demonstrate that AZD2858 can markedly enhance the cytotoxic and cytostatic effects of SN-38 and FOLFIRI through a combination of increased apoptosis and growth arrest according to dosage and treatment conditions. Figure 6 extends this analysis to cells cultured as spheroids, sometimes considered to better represent tumor responses compared to single cell cultures.

### Major comments

Most of the data presented is of good technical quality and supports the conclusions drawn. There are however a small number of instances where this is not true; ie where the data are of insufficient technical quality, or where the description or interpretation of the results is at variance with the data which is presented. Some examples:

1) Fig.2E - the claim that “we observed an increase in RPA, Topb1 and Pol-epsilon levels when CPT and AZD2858 were added together” do not seem to be justified by the data provided. It is also unclear what the purpose/ significance of this experiment is.

Thank you for pointing out the contradiction in Figure 2E. Upon review, we identified an error in the labeling of conditions (CPT and AZD2858 were inadvertently swapped). The corrected figure now clearly shows that, at the 60-minute timepoint after starting replication, the combination of CPT and AZD2858 results in a greater accumulation of TopBP1, Pol ε, and RPA on chromatin compared to CPT alone. We have revised the sentence to: “Our data demonstrate that combining CPT and AZD2858 earlier enhances the accumulation of replication-related factors (RPA, TopBP1, and Pol ε) on chromatin compared to CPT treatment alone, particularly visible at the 60-minute after starting replication.”

The significance of this experiment lies in its connection to the earlier observation that AZD2858 restores BrdU incorporation when combined with SN-38, as shown in flow cytometry data (Figure 3A). At a molecular level, this was further supported by DNA fiber assays, which revealed that replication tracks (CldU tracts) were longer in the combination treatment compared to SN-38 alone (Figure 3B).

To strengthen and validate these findings, we chose to employ the Xenopus egg extract system for several reasons. This model provides a highly controlled environment where DNA replication occurs without confounding effects from transcription or translation. Moreover, replication is limited to a single round, offering a unique opportunity to specifically interrogate replication mechanisms. These attributes make the Xenopus model an ideal system to confirm that AZD2858 facilitates replication recovery in the presence of replication stress induced by agents like CPT. This will lead, in longer treatment, to accumulation of DNA damage and apoptosis (Figure 3D-E and Figure 4A-D)

2) Figs. 3 A and C certainly show that the SN-38-mediated suppression of DNA synthesis is modified and partially alleviated by co-treatment with AZD2858. The statement however that “prolonged co-incubation with AZD2858 for 6 and 12 hours effectively abolished the SN-38 induced S-phase checkpoint” is clearly misleading. If this were true, then the BrdU incorporation profiles of the respective samples would be similar or identical to control, which clearly they are not. Clearly AZD2858 is affecting the imposition of the S-phase checkpoint in some way, but not “abolishing” it.

We appreciate the reviewer’s detailed observations regarding Figures 3A and **3C** and the phrasing in our manuscript. We agree that the term “abolished” is not precise in describing the effects of AZD2858 on the SN-38-induced S-phase checkpoint.

To clarify: our data indicate that co-treatment with AZD2858 modifies and partially alleviates the SN-38-induced suppression of DNA synthesis, as demonstrated by increased BrdU incorporation relative to SN-38 treatment alone. However, as the reviewer correctly points out, the BrdU incorporation profiles of the co-treated samples do not fully return to control non treated cells levels. This suggests that while AZD2858 significantly mitigates the S-phase checkpoint, it does not completely abolish it.

We have revised the statement in the manuscript to better reflect these findings, as follows: “Prolonged co-incubation with AZD2858 for 6 and 12 hours significantly alleviated the SN-38-induced S-phase checkpoint, as evidenced by the partially increased BrdU incorporation. However, the population of co-treated cells is heterogeneous: some cells exhibit BrdU incorporation levels similar to those of untreated control cells, while others incorporate BrdU at levels comparable to cells treated with SN-38 alone. This indicates that AZD2858 does not fully restore DNA synthesis to control levels across the entire cell population.”

This revised phrasing aligns with the data presented and acknowledges the partial recovery of DNA synthesis observed. Thank you for bringing this to our attention and helping us improve the accuracy of our conclusions.

3) Fig. 3 E. The western blots of pDNA-PKcs (S2056) and total DNA-PKcs are really not interpretable. It is possible to sympathise that these reagents are probably extremely difficult to work with and obtain clear results, however uninterpretable results are not acceptable.

We agree that the data presented in the Fig3E are difficult to interpret. As noted by Reviewer 1, we recognize the challenge of obtaining clear and reliable results with these specific reagents. Based on this feedback, and to ensure the robustness of our conclusions, we have decided to exclude these specifics blots from the revised manuscript.

We believe that this adjustment will enhance the clarity and reliability of the manuscript while focusing on the other, more interpretable data presented. Thank you for pointing this out, and we appreciate your understanding.

4) Fig. 3D. This is a puzzling image. Described as a PFGE assay, it presumably depicts an agarose gel, with intact genomic DNA at the top and a discrete band below representing fragmented genomic DNA. This is a little surprising, as fragmented genomic DNA does not usually appear as a specific band but as a heterogenous population or “smear”. Nevertheless, even if one accepts this premise, it is unclear what is meant by “DSBs remained elevated after the combined treatment” when the intensity of this band is equivalent for both SN-38 and SN-38 + AZD2858 treatments.

We thank the reviewer for his insightful comments regarding the PFGE results in Figure 3D. We agree that the appearance of a discrete band, rather than a heterogeneous smear, is atypical for fragmented genomic DNA in this assay. However, by enhancing the signal intensity (as shown below), the expected smear becomes more appreciable.

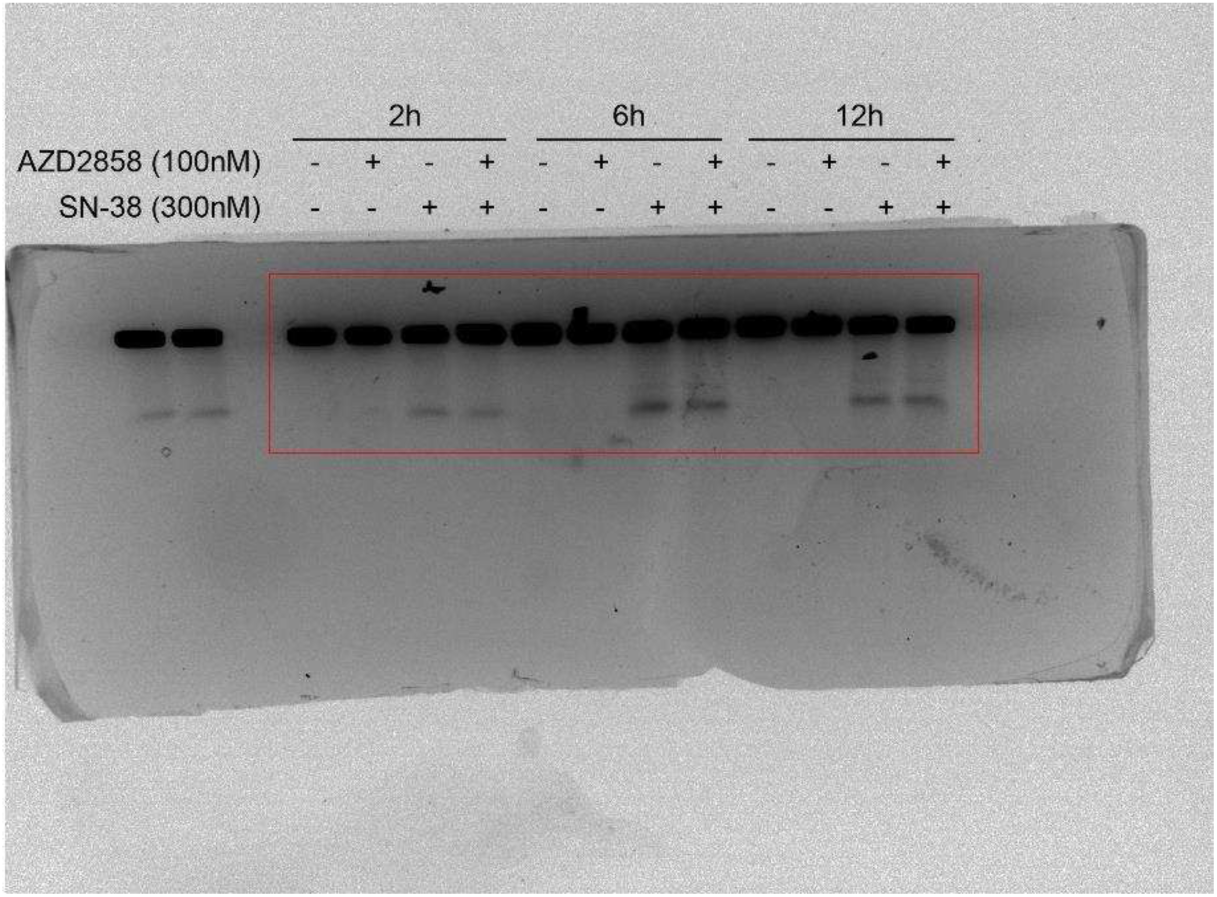

Regarding the interpretation of the band intensities, we agree that the signals for SN-38 and SN-38 + AZD2858 appear similar under these specific conditions. At the relatively high concentration of SN-38 used in this experiment (300 nM), it is indeed challenging to observe a more pronounced effect on DNA breaks. This is why we proposed the “DSBs remained elevated after the combined treatment” because the band intensity of SN-38 single agent treated cells or combined with AZD2858 is comparable. However, we note a slightly more intense γH2AX signal over time when AZD2858 is combined with SN-38 compared to SN-38 alone (Figure 3E). Furthermore, under lower, sub-optimal doses of SN-38 and over extended incubation treatment (48h), we observe a clearer increase in fragmented DNA bands, as demonstrated in Figure 4D.

### Minor comments

1) Fig. 1. A surprisingly large number of compounds scored positive in the primary screen for inhibition of Topbp1 condensation (>300). Of the 131 of these selected for secondary screening using Chk1 activation (S345 phosphorylation) as a readout approximately 2/3 were negative, implying that a majority of the tested compounds inhibited Topbp1 condensation but not Chk1 activation. What could explain that?

Thank you for this thoughtful comment. The discrepancy between the large number of compounds scoring positive for TopBP1 condensation inhibition and the smaller number inhibiting Chk1 activation (S345 phosphorylation) could be attributed to several factors:

- Different cell lines and induction methods: The initial screen was conducted in HEK293 Trex-Flpin cells overexpressing optoTopBP1, while the secondary screen used HCT116 cells. In addition, the methods used to induce the respective pathways were distinct: in the primary screen, we employed a blue light induction of opto-TopBP1 condensates, whereas in the secondary screen, we used an SN-38 treatment to induce DNA replication stress and activate the Chk1 pathway. These differences could account for the varying responses observed in the two screens.
- The compounds that inhibited TopBP1 condensation might not fully block Chk1 activation. While they disrupt TopBP1 condensation, they may still allow for partial activation of Chk1 or Chk1 activation through alternative mechanisms. For instance, Chk1 activation could be mediated by other signaling pathways or molecules, such as ETAA1, a known Chk1 activator (1). Thus, TopBP1 condensation inhibition does not necessarily translate to complete inhibition of Chk1 activation, especially if ETAA1 is employed by cells as a rescue activator.
- Some compounds may affect chromosome dynamics, potentially generating mechanical forces or torsional stress that could activate the ATR/Chk1 pathway independently of TopBP1 (2).

These factors suggest that while the compounds effectively disrupt TopBP1 condensation, they may not always fully inhibit the downstream Chk1 activation, pointing to the complexity of the DNA damage response pathways.

2) Fig. 2D. The protein-protein interaction assay shown demonstrates that AZD2858 ablates the light-induced auto-interaction between exogenous opto-Topbp1 molecules and ATR plus or minus SN-38, but clearly endogenous Topbp1 molecules do not participate. Why is this?

The biotin proximity labeling assay was conducted without exposing cells to light, using a TurboID module fused to TopBP1-mCherry-CRY2. Stable cell lines were then generated in HEK293 Trex-FlpIn cells, where endogenous TopBP1 is still expressed. Upon adding doxycycline, the recombinant TurboID-TopBP1-mCherry-Cry2 (opto-TopBP1) is induced at levels comparable to endogenous TopBP1 (Fig 2D).

Since the opto-TopBP1 construct exhibits behavior similar to that of endogenous TopBP1 (1), we used it to investigate whether TopBP1 self-assembly and its interaction with ATR are influenced by AZD2858 alone or in combination with SN38. Our results show that treatment with SN38 increases the proximity between opto-TopBP1 and the endogenous TopBP1 (not fused to TurboID). However, AZD2858, either alone or in combination with SN38, disrupts the self-assembly of recombinant TopBP1 with itself as well as its interaction with endogenous TopBP1.

Reviewer #3 (Significance (Required)):

### Significance

Liquid phase separation of protein complexes is increasingly recognised as a fundamental mechanism in signal transduction and other cellular processes. One recent and important example was that of Topbp1, whose condensation in response to DNA damage is required for efficient activation of the ATR-Chk1 pathway. The current study asks a related but distinct question; can protein condensation be targeted by drugs to manipulate signalling pathways which in the main rely on protein kinase cascades?

Here, the authors identify an inhibitor of GSK3-beta as a novel inhibitor of DNA damage-induced Topbp1 condensation and thus of ATR-Chk1 signalling.

This work will be of interest to researchers in the fields of DNA damage signalling, biophysics of protein condensation, and cancer chemotherapy.

